# Are trapping data still suited for home range estimation? An analysis with various estimators, asymptotic models and data ordering procedures

**DOI:** 10.1101/2022.03.15.484432

**Authors:** L. Socias-Martínez, L. R. Peckre, M. J. Noonan

## Abstract

Understanding the size of animals’ home ranges is vital for studies in ecology and conservation. Trapping datasets are an important source of information when targeting the biodiversity of an area, inconspicuous species, or high numbers of individuals in contrast to more expensive telemetry-based methods such as radio- or GPS-collaring. Currently, studies relying on trapping lack an evaluation of the performance of existing home range estimation procedures comparable to those developed for telemetry. Using animal movement simulations, we evaluate three variables reflecting the trade-offs faced by ecologists when designing a trapping study, 1) the number of observations obtained per individual, 2) the trap density and 3) the proportion of the home range area falling inside of the trapping grid. We compare the performance of five estimators on these conditions, four commonly used (AKDE, KDE, MCP, LoCoH) and a possible alternative for situations with low trap density or high number of observations (bicubic interpolation). We further test suggested benefits of using asymptotic models (Michaelis-Menten and monomolecular) to assess the total home range area when information obtained per individual is scarce, as this situation might be common in trapping datasets. In addition, we propose sorting the observations based on the distance between locations to improve the performance of asymptotic models’ estimates. Using the results of the different procedures we constructed a generalized additive model (GAM) that allows predicting the bias in home range size under the different scenarios investigated. Our results show that the proportion of the area covered by the trapping grid and the number of observations were the most important factors predicting the accuracy and reliability of the estimates. The use of asymptotic models helped obtaining an accurate estimation at lower sample sizes and this effect was further improved by distance-ordering. The autocorrelation informed KDE was the estimator performing best under most conditions evaluated. Nevertheless, bicubic interpolation can be an alternative under common trapping conditions with low density of traps and low area covered. We provide the current results to the constructed GAM as a prospective tool for ecologists planning a new study or with already collected datasets that aim at assessing the potential biases in their estimates. Reliable and accurate home range estimates using trapping data can optimize monetary costs of home range studies, potentially enlarging the span of species, researchers and questions studied in ecology and conservation.

## Introduction

Home ranges have been defined as “…the area traversed by the individual in its normal activities of food gathering, mating, and caring for young” (Burt 1943). The intensity of resource use and competition, the amount of gene flow, the mating system and social structure, and the spread of information and diseases might all be inferred through animals’ space use (Mueller and Fagan 2008, Schick et al. 2008, Dammhahn and Kappeler 2008, Nathan et al. 2008). Knowledge on a species’ use of space is therefore at the core of understanding biodiversity and design effective conservation measures (Law and Dickman 1998, Allen and Singh 2016, Oppel et al. 2018).

Although the ecological definition of the home range is not specific to any data type nor estimator, the accurate estimation of home ranges is challenged by mismatches between the methods’ assumptions and the nature of the data collected (Hayne 1949, Andrzejewski 2002, Powell and Mitchell 2012, Noonan et al. 2019, Wszola et al. 2019, Fleming et al. 2019). During the last century, an extensive body of literature has been directed to describe and quantify biases arising from methods used (see e.g., Hayne 1949, 1950, Worton 1989, Harris et al. 1990, Seaman et al. 1999, Andrzejewski 2002, Laver and Kelly 2008, Fleming et al. 2018, 2019). Comparisons of different estimators for home ranges have suggested strengths of each under different conditions (e.g., sample size, the time between measurements and autocorrelation) (Hayne 1949, Van Winkle 1975, Rose 1982, Worton 1987, Plotz et al. 2016, Halbrook and Petach 2018, Noonan et al. 2019, Vieira et al. 2019). Further spatial and temporal heterogeneities have also been investigated (e.g., ecotypes, physical boundaries) and shown to have an effect (Ouellette and Cardille 2011, Halbrook and Petach 2018, Wszola et al. 2019). The (mis)match between weaknesses and strengths of estimators and data collected is therefore a major potential source of error in home range estimates that needs to be evaluated (Hayne 1949, Van Winkle 1975, Worton 1989, Seaman et al. 1999, Kie et al. 2010, Powell and Mitchell 2012, Fleming and Calabrese 2017, Halbrook and Petach 2018, Fleming et al. 2019).

Several authors have suggested that if animals exhibit site fidelity (e.g. Ebersole, 1980; Halbrook & Petach, 2018; Heupel, Simpfendorfer, & Hueter, 2004; Powell, Zimmerman, & Seaman, 1997; Reid & Weatherhead, 1988; Spencer, Cameron, & Swihart, 1990), the information on space use obtained should follow a saturation curve (Harris et al. 1990, Haines et al. 2009, Soanes et al. 2013, Leo et al. 2016, Halbrook and Petach 2018, Wszola et al. 2019). Deviations from this saturation may thus indicate that i) the animals being monitored are not range-resident (e.g., (Morato et al. 2016)), or ii) the individual has been monitored for too short of a period for the range-resident behavior to be observable (Fleming et al. 2014). The latter problem represents a key challenge that can bias home range estimates and any subsequent hypothesis testing, or decision making. Modelling the saturation curve can allow estimating the true home range even if the information is still incomplete (e.g., (Soanes et al. 2013, Leo et al. 2016, Halbrook and Petach 2018)), thus offering the possibility of reducing the amount of data needed for accurate estimates. Nevertheless, most empirical studies do not make use of asymptotic models perhaps because a thorough examination of the performance of this type of analyses is lacking.

The mentioned limitations and mismatches between the methods used for obtention of information and description of space use in animals might be even more pronounced in trapping-based studies. While trapping studies were once predominant in home range estimation (Mohr 1947, Hayne 1949, Andrzejewski 2002), they have decayed in the last decades in favor of newer, more accurate and reliable technologies based on radiotracking or GPS (Innes and Skipworth 1983, Ward 1984, Bergstrom 1988, Ribble et al. 2002, Gil-Sánchez et al. 2011, Kays et al. 2015). Trapping studies suffer most importantly from extreme uncertainty on the information obtained per individual which usually consists of few observations (Dammhahn and Kappeler 2005, Lira and Fernandez 2009, Gil-Sánchez et al. 2011, Kane et al. 2015, Kumbhojkar et al. 2020). Unlike for telemetry data, where temporal autocorrelation is the predominate source of bias for home range estimation (Noonan et al. 2019, Silva et al. 2022), small sample sizes are a key consideration. All home range estimators are sensitive to sample size (Schoener 1981, Fleming et al. 2019). Thus, the reliability of information on home ranges coming from trapping studies is often uncertain and an evaluation of the necessary conditions for accuracy and reliability are needed.

In addition to the low number of observations, the size of the trapping grid and its emplacement relative to the center of an individual’s home range can also impact the measurements in trapping studies (Rajska-Jurgiel 2001, Lukacs et al. 2005, Bondrup-Nielsen 2011, Sun et al. 2014). Due to a lack of understanding on the biases and limitations of carrying home range analyses with trapping data, researchers with trapping datasets face extreme uncertainty on the error of their estimates and might even dismiss conducting home range analyses altogether (Fig. 1). This situation can bias the knowledge on movement ecology to only those species that can be monitored via telemetry. Trapping data harbor enormous potential for home range analysis due to the range of species that can be studied and the reduced budgetary constraints. For instance, camera- or live-trapping can be used to study animals that are too small to carry a telemetry or GPS collar. Moreover, with the average all-in cost of tracking an animal being ca. $10,000 USD (Thomas et al. 2011), collars are expensive, reducing the number of individuals and species that can be monitored at a given location. In addition, the costs of telemetry-based methods might set a boundary to the economic background of institutions or researchers in movement ecology. If trapping based studies prove more reliable than previously thought they could enlarge the scope of researchers, species and time of monitoring helping improve general knowledge on movement ecology essential for fundamental and applied science.

**Figure 1:**
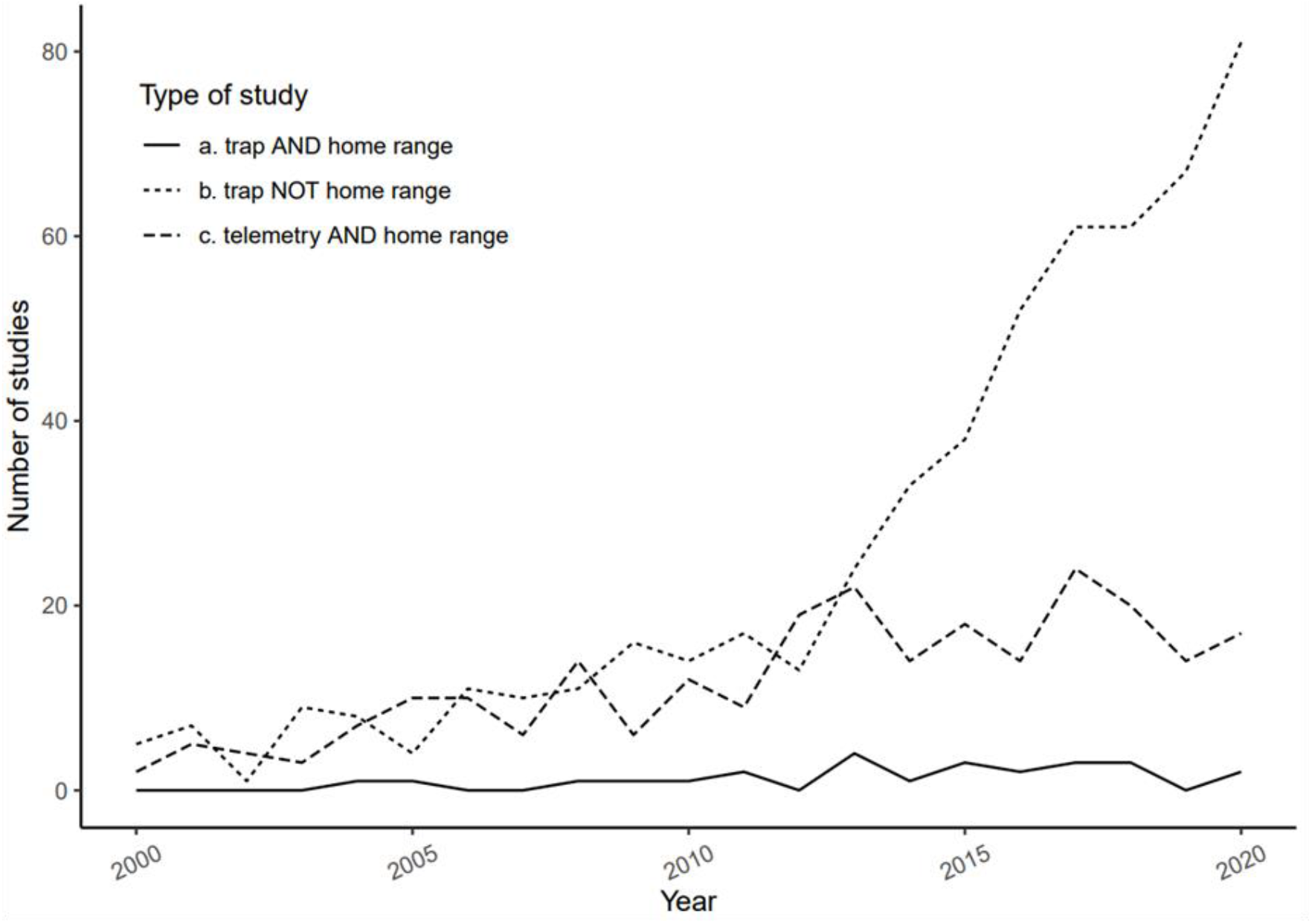
Published studies using telemetry and trapping data: Results from a search in ScienceDirect for article sections title-abstract-keywords using the search words: a. ‘ (“camera trap” OR “live trap”) AND “home range” ‘ for studies using trapping datasets evaluating home ranges, b. ‘ (“camera trap” OR “live trap”) NOT “home range” ‘ for studies collecting trapping datasets but not using them for home ranges and c. ‘(GPS OR radio) AND “home range” ‘ for studies on home range using telemetry methods. Only articles in the fields “Agricultural and Biological Sciences”, “Environmental Science” and “Biochemistry, Genetics and Molecular Biology” were included.

To fill the current gap in knowledge on trapping datasets for home range analyses we conducted a thorough evaluation of their accuracy under a vast array of conditions in a set of simulations. We evaluated the performance of five estimators, two kernel based (autocorrelated kernel density estimation AKDE, and traditional KDE), a polygon-based (minimum convex polygon MCP) and its generalization (local convex hull LoCoH) and an interpolation-based method (bicubic interpolation BicubIt). Moreover, we evaluated the performance of two asymptotic model predictions (Michaelis-Menten MicMen and Monomolecular MonMol) as they have been suggested to help when information is incomplete, a common situation for trapping datasets. In addition, we evaluated these asymptotic models on two conditions for ordering the observations, time-ordered and distance-ordered as we suspected that the latter could improve their predictions. The combinations of all mentioned variables were evaluated under three key parameters for designing a trapping study: the trap density, the effective sample size, and the proportion of the home range covered by the trapping grid. These measures correspond to the amount of effort and money invested in a study based on animal captures and can impact the estimates of home range size. Our analyses thus may point to optimal investments for describing animal home ranges in studies relying on animal trapping, representing an important tool for ecologists that aim at study movement using trap data.

## Material and Methods

### Simulating animal movements and trapping

We simulated home ranges with a circular area of 100 Hectares and a uniform distribution probability for observing the animal across it. We simulated movement for 100 individuals moving within this type of home range without any autocorrelation (i.e., using a I.I.D. movement model), as trap data are likely to be free from meaningful autocorrelation. To capture the individuals, we generated a series of trapping grids with varying densities measured as number of traps in each line of the grid per home range radius (ranging from 2 to 42 over intervals of 10, see Fig. 2). Each trapping grid was placed with its center at different distances from the center of the home range creating a coverage ranging from 20 to 100% (see Fig. 2). We considered an animal captured if an observation fell within 200m of a trap (approx. corresponds to 20m for small animals with home ranges of 1Ha). We chose this measure to simulate the attraction of baiting in real-world scenarios with live trapping but can also be understood as a metric of visibility for camera traps. Finally, for each individual in each condition of trap density and area covered we simulated positions until it was captured 2000 times. The scripts necessary to reproduce these simulations will be provided when published in a peer-review journal.

**Figure 2:**
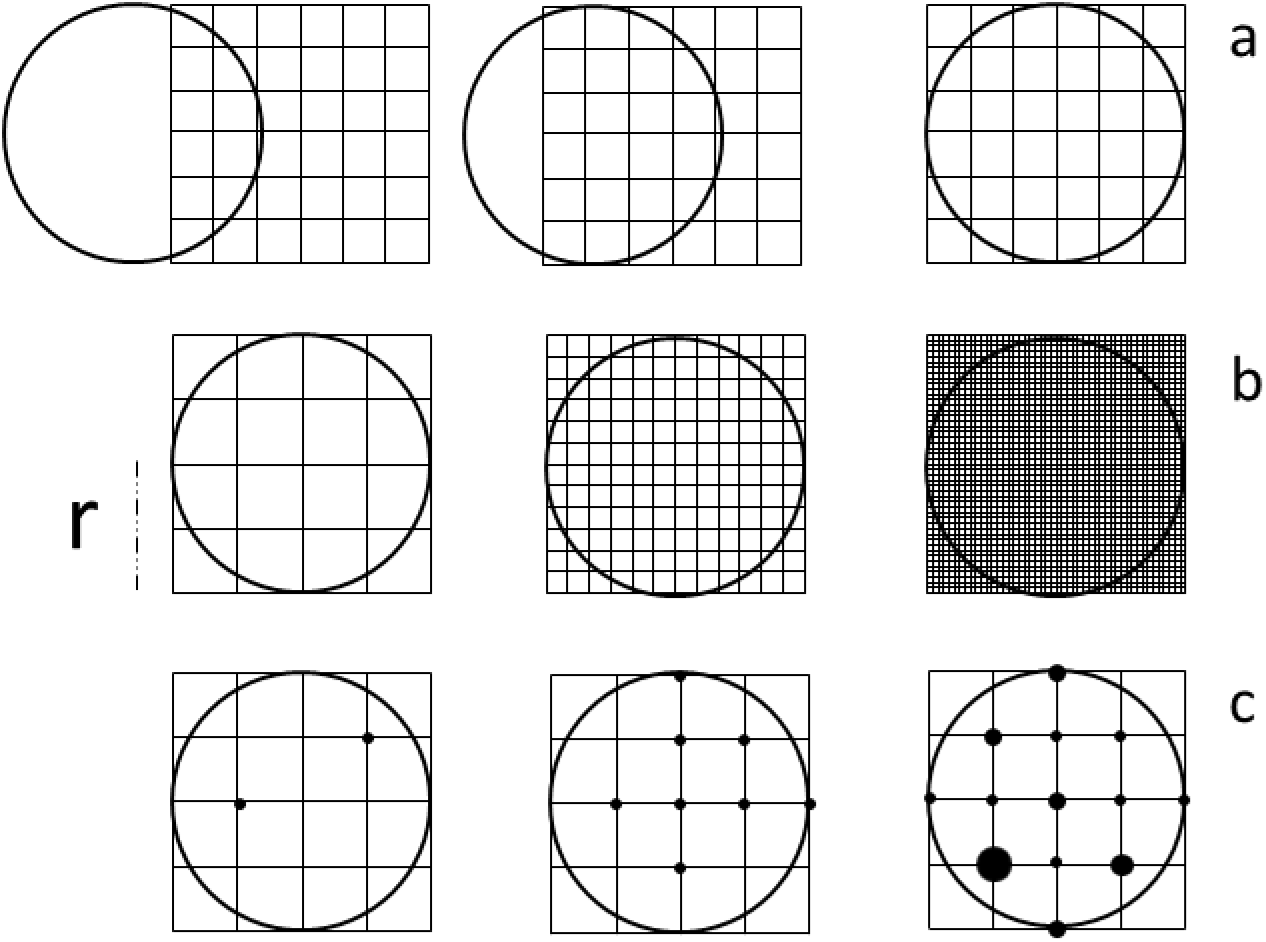
The three variables of interest for study design: **a**. The area covered by the trapping grid, with low, medium and total coverage from left to right, **b**. the trap density, with again low, medium and high values of traps per home range radius (illustrated with the dashed bar and the “r” symbol), and **c**. the number of observations, with dots indicating their location and the size of dots their number in a given location.

### Fitting estimators and asymptotic models

#### Incremental analyses: home range area as a function of the number of observations

For each dataset we decided a set of ticks between 2 and 2000 observations on which to conduct the estimates of home ranges. The interval between ticks increased progressively as we expected redundancy in information because of the asymptotic behavior of spatial variance for range-resident animals. Based on these observation number ticks we performed two orderings of the locations for each individual. The first corresponds to standard use in home range analyses, with observations ordered by time (time-ordered hereafter). The second one was achieved by taking each subset of observations from the time ordered dataset and reordering based on distance using farthest point sampling (Fig. 3). Farthest point sampling allows reordering the data, with each point being followed by its corresponding farthest point in the dataset based on Euclidean distances, excluding computed points already used. By doing so, while for each number of observations both the time ordered and the distance ordered dataset contained the same exact records of space positions, the distance ordered included most information on space use within the first observations (Fig. 3).

**Figure 3:**
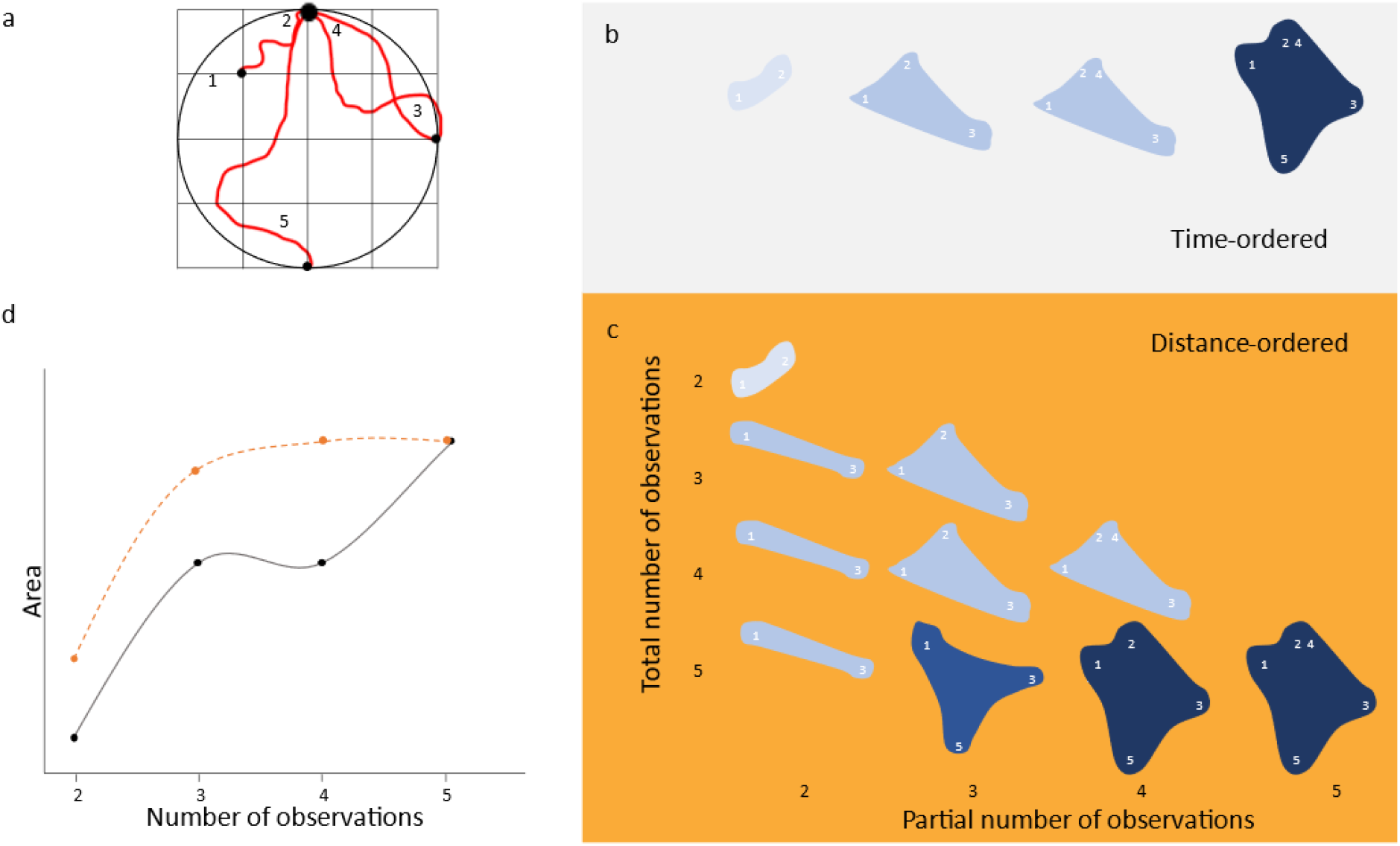
Time-ordered and distance-ordered sampling curves: **a**. A hypothetical trapping of an individual with 5 observations, **b**. the succession of home range areas calculated from a. with standard time-ordered incremental analyses, **c**. the different areas estimated from a. by ordering the data based on farthest-point sampling with partial datasets from two to five observations and **d**. representation of the different curves obtained with five observations using either time-ordered (grey continuous) or distance-ordered (orange dashed) incremental analyses.

For each dataset we calculated the home range area incrementally. For the time-ordered dataset this incremental analysis corresponded to calculate the home range area at each tick (Area_i_), yielding a unique incrementing curve of home range area per dataset from 2 to 2000 observations. For the distance-ordered dataset a different curve ending at each tick was calculated (Fig. 3). Each tick had a different curve because each subset had different farthest points being redistributed accordingly with more information clumping in the first observations.

#### Home range estimators

To calculate the home range areas, five estimators were used. A widely used method, the Kernel Density Estimator (KDE) was chosen as it is the most statistically efficient non-parametric density estimator. We further used a refined version of the KDE that takes autocorrelation into account (AKDE) and has further refinements for small sample sizes, irregular sampling, and measurement error (Silva et al. 2022). For KDE, an automatic bandwidth selection method was applied to each subset of the data based on bisection algorithm that finds the smallest bandwidth generating *n* polygons (Kie 2013). We set *n* to 1 to have a single continuous home range area. Two commonly used polygon-based methods, the minimum convex polygon (MCP) and its generalization, the local convex hull (LoCoh) were also used (see Supporting information, SI). Furthermore, we explored the properties of a home range estimation based on bicubic interpolation (BicubIt). We expect this latter estimator to perform better under low trap densities as well as with very large datasets when kernel-based estimators become ineffective due to the bandwidth limiting to 0 when *n*→∞(see SI).

#### Asymptotic models

A Michaelis-Menten model as described in (Leo et al. 2016) (eq 1) was fitted to the home range area as a function of the number of observations with automatic initial values for the rest of parameters. The Michaelis-Menten model is a saturating model that is well known in the biological sciences. A challenge with this model, however, is that the asymptote can be difficult to estimate with data (Bolker 2008). We therefore also explored the potential benefits of the monomolecular model (eq 2). The models were fitted to each tick in the single curve for the time-ordered dataset using Levenberg-Marquardt nonlinear least-squares algorithm. For the distance-ordered, the models were fitted to each curve created with the observations until a given tick. The predicted asymptotes were retrieved and assigned as value for the home range area to the number of observations it used.

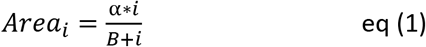

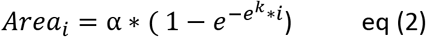

### Evaluating the different methods for home range estimation on trapping data

We transformed the areas obtained in the previous analyses to a proportion of the true 95% home-range area. Using the variables describing the estimator used, the time-ordering procedure, the asymptotic model used and the three variables for study design (number of observations, trap density and area covered) we modelled the accuracy of area estimates using a generalized additive model (GAM) in the mgcv package (Wood 2017) in R version 4.0.5 (R Core Team 2019). GAM was chosen to capture the possible complex non-linear relationships between our variables of interest and the home range area predicted. The proportion of the true area was made dependent on the number of observations, the trap density and the area covered by the trapping grid using a gaussian link function (eq. 3). Since asymptotic models can overestimate the area dramatically under low number of observations, we filtered out extreme outliers before fitting the GAM. We defined extreme outliers as those that overestimated or underestimated the true area by a factor of 10. This type of aberrant predictions can be easily detected by researchers in their datasets once some knowledge on the species is available.

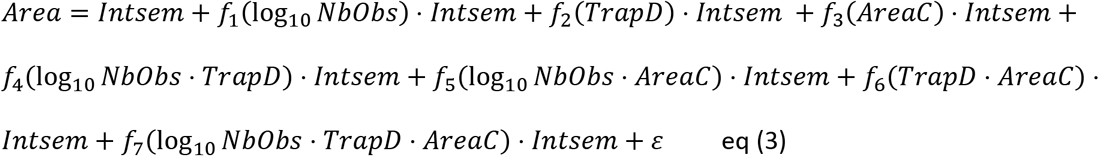

*where f*_1−3_ *are smooth and f*_4−7_ *partial tensor product interaction smooth functions estimated by restricted maximum likelihood*

The number of observations was transformed using its base ten logarithm and the knots for the basis functions were placed according to a logarithm function in the range of observed values. Placing the knots in this way allowed to model the variation at low observation numbers when small changes are likely to induce much stronger effects than at the higher end. We included a three-way interaction as a tensor product interaction between the number of observations, the trap density and the area covered as their effects are all conditional on each other (eq. 3). All lower-level interactions combinations and main effects of the three variables were also included as tensor product interactions and smooth terms respectively.

Furthermore, a factor variable gathering the home range estimator, the asymptotic model and the sampling ordering procedure (intsem) was generated. “intsem” was included as a main effect and as a “by variable” on each smooth and tensor product interaction term. Including this smooth-by-factor interactions allowed fitting a smooth term separately for each level of “intsem”. Such flexibility was needed to capture the likely different shapes that the functions for each level and combinations of the variables of interest might have. The use of a common variable, “intsem”, reduced the computational burden of modelling the home range estimator, the asymptotic model and the sampling order procedure as separate terms. The sufficiency of the basis dimensions for each smooth term were evaluated by using the standard procedures for GAM evaluation in the mgcv package including the tests of gam.check function and the diagnostics of linear modelling assumptions. For further details on other parameters see SM.

We then evaluated the accuracy and reliability of home range estimates in trapping settings using our simulations. To visually identify the parameter spaces of interest for trapping studies, predictions were considered accurate if fell within ±10% of the true area and reliable if the confidence interval had a breadth not exceeding ±20% of the estimate. For each graphic representation we described with emphasis only the values that were accurate and reliable according to the above definitions. When possible, we also described the conditions that allowed reaching accuracy instead of under- or over-estimation and reliability.

To produce the graphic representations summarizing the results, we predicted the home range area as a function of the two-way interactions between the number of observations and the trapping density (while setting the area covered to 100%), between the number of observations and the area covered (setting the trap density to 2), and between the area covered and the trap density (setting the number of observations to 7) using the constructed GAM. The fixed values of low trap density and low number of observations were chosen to describe possible common situations in field studies using trapping procedures. Further graphic representations including fixed values at low, mean, and high values can be found in the SI.

## Results

The 43 individuals simulated yielded a total number of 415,612 observations based on the combination of 5 estimators, 2 ordering procedures, 3 model types (including no model), 6 trap densities, 6 area covered proportions and 28 different number of observations. Overall, the GAM model performed satisfactorily with most residuals being close to 0 (see SI). There existed some level of heteroskedasticity with high variance characterizing the areas at low number of observations. Moreover, residuals were non-normally distributed due to a positive skew resulting from i) the distribution of the response (bounded at 0 but unbounded at positive infinity) and ii) high overestimated asymptotic models’ predictions at low number of observations (see SI).

### Interacting effects of number of observations and trap density

When the trapping grid covered the home range entirely, the number of observations was the primary factor driving the accuracy of the estimates (Fig. 4 see vertical color bands and accuracy niches). The reliability of estimates also depended more strongly on the number of observations than on the trap density with an estimator-specific minimum above which estimates were consistently (Fig. 4). The estimators that conformed more strongly to the described general patterns were the two Kernel-based. The two polygon-based estimators followed in conformity, deviating from the described pattern when used with time-ordered asymptotic models where accuracy niches reacted more strongly to trap density.

**Figure 4:**
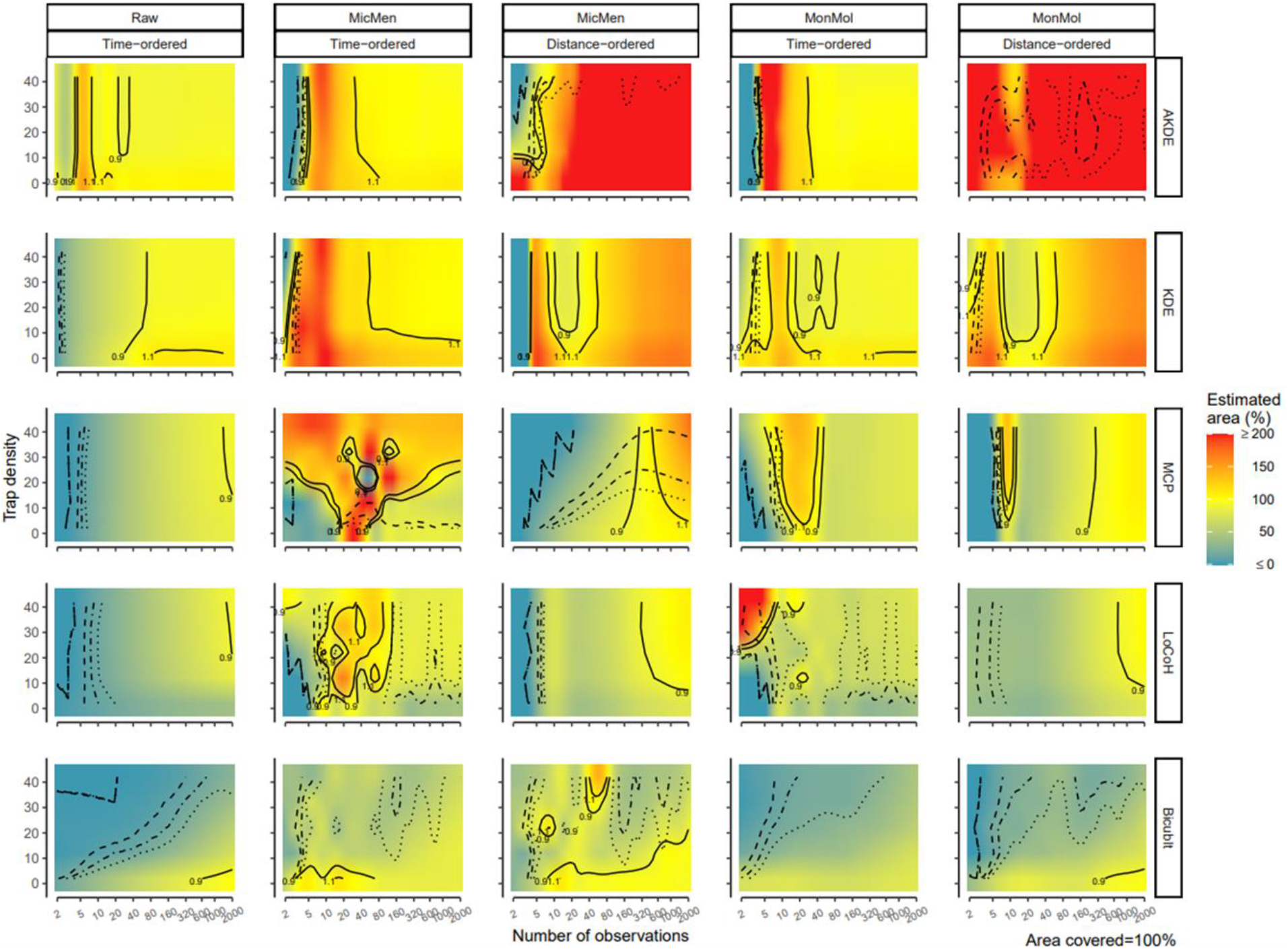
Interacting effects of number of observations and trap density on home range area estimations: Rows gather different estimators, from top: autocorrelated Kernel density estimator (AKDE), the traditional version (KDE), minimum convex polygon (MCP), local convex hull (LoCoH) and bicubic interpolation (Bicublt). Columns gather the combination of two factors, the ordering procedure (time or distance) and the asymptotic model used (“Raw” indicating no model, “MicMen” Michaelis-Menten and “MonMol” Monomolecular). The x axis displays The number of observations and the y axis the trap density, the area covered was fixed at 100%. The colors in the plots indicate the estimated area as a percentage of the true area predicted by the GAM model, yellow values approximate the true area, and red indicate over-estimation while blue underestimation. Values above 200% where trimmed and given a 200 while values below 0% where also trimmed as 0 to concentrate color variation on a meaningful range. The solid lines indicate the contour of accurate estimations, defined as ±10% of the true area. Pointed lines indicate the reliable estimations measured as those where the confidence interval was ±20% of the estimated area, point-dashed ±40% and dashed ±60%. Finally, on the lower-right corner the fixed variable value is indicated.

BicubIt stand out from the rest by having the accuracy and reliability dependent equally on the number of observations and the trap density (Fig. 4 see diagonal color bands and lines).

The two Kernel density estimators outperformed the rest with accurate estimates at 9 and 20 observations (AKDE and KDE respectively). The rest underestimated the true area before ∼1000 (MCP and LoCoH) and ∼600 (BicubIt, only at low trap density) observations where gathered. The reliability of estimates was obtained at very low number of observations except for BicubIt where the higher the trap density the more observations were needed for a reliable estimate.

Interestingly, using the asymptotic models generated accuracy and reliability patterns that had estimator-specific optima generally lacking asymptotic consistency (i.e., adding more observations after the accuracy and reliability niche was attained induced its loss). The two Kernel-based methods suffered by needing more observations for an accurate estimate than when using the raw data alone, particularly at low trap densities. At higher trap densities, though, KDE benefited from using MonMol through improved accuracy of estimates at low sample sizes. MCP became unreliable when used with MicMen and never reached accuracy while MonMol allowed accurate and reliable estimates at lower number of observations. LoCoH benefitted from using MicMen by reaching reliability and accuracy at ∼10 observations while using MonMol led to very limited accuracy niches. Similarly, BicubIt benefited only from MicMen and not from MonMol. MicMen allowed to have accurate and reliable predictions between 3 and 80 observations for the smallest trap densities.

Distance-ordering helped through stabilization of the accuracy and reliability niches after their advent and through a higher set of conditions yielding reliable estimates. Nevertheless, this procedure only benefited accuracy with KDE by reducing the minimum number of observations and with BicubIt by expanding the range of trap densities yielding an accurate estimate.

### Interacting effects of number of observations and area covered

When the trap density was four traps per grid line per home range radius, increasing both observations and area covered led to higher accuracy. Reliability depended on an estimator-specific minimum number of observations above which increasing the area covered led to more reliable estimates. The accuracy of Kernel-based estimators was affected more strongly by the area covered than by the number of observations (Fig. 5, see “v” shaped accuracy niches along the x axis). The polygon-based estimators were, on the contrary, more affected by the number of observations, once a minimum area covered was attained. BicubIt offered a bimodal accuracy were the extremes of area covered tended to yield more accurate estimates. AKDE performed best, followed by KDE, BicubIt, MCP and LoCoH. AKDE was able to offer accurate and reliable estimates with 5 observations and an area covered as low as 50%, while KDE needed at least 40 observations. MCP and LoCoH were unable to offer accurate estimates while BicubIt needed the highest area covered and number of observations.

**Figure 5:**
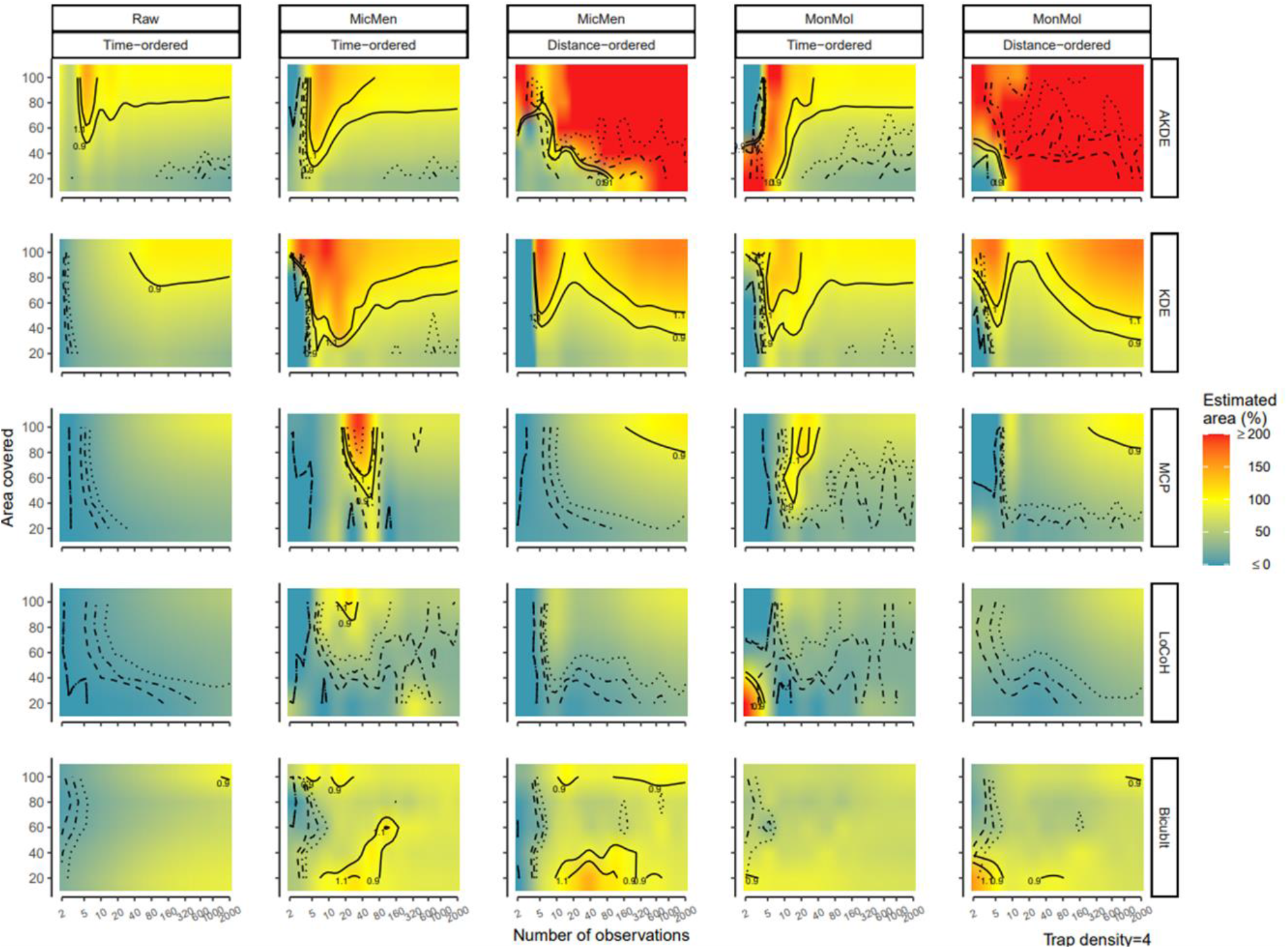
Interacting effects of number of observations and area covered by the grid on home range area estimations: Rows gather different estimators, from top: autocorrelated Kernel density estimator (AKDE), the traditional version (KDE), minimum convex polygon (MCP), local convex hull (LoCoH) and bicubic interpolation (Bicublt). Columns gather the combination of two factors, the ordering procedure (time or distance) and the asymptotic model used (“Raw” indicating no model, “MicMen” Michaelis-Menten and “MonMol” Monomolecular). The x axis displays the number of observations and the y axis the proportion of the area covered by the trapping grid, the trap density was fixed at 4 traps per HR radius. The colors in the plots indicate the estimated area as a percentage of the true area predicted by the GAM model, yellow values approximate the true area, and red indicate over-estimation while blue underestimation. Values above 200% where trimmed and given a 200 while values below 0% where also trimmed as 0 to concentrate color variation on a meaningful range. The solid lines indicate the contour of accurate estimations, defined as ±10% of the true area. Pointed lines indicate the reliable estimations measured as those where the confidence interval was ±20% of the estimated area, point-dashed ±40% and dashed ±60%. Finally, on the lower-right corner the fixed variable value is indicated.

Using asymptotic models offered some benefits albeit the accuracy niches were not consistent when increasing sample sizes. Accuracy niches had a “v” shape (except for BicubIt, indicating optimal values changing with the upper space in the “v” risking overestimation. AKDE and KDE benefited by lowering the minimum area covered (∼30%) in order to retrieve an accurate estimation. MCP benefited from MonMol reducing the minimum number of observations to∼15, while MicMen yielded unreliable estimates. Both asymptotic models helped LoCoH achieve accurate estimates, despite only MicMen with reliability at ∼15 observations. BicubIt benefited from asymptotic models from increased reliability and accuracy, but more so from MicMen. Using MicMen allowed to retrieve accurate estimations at as low as 4-5 observations with both high and low area covered.

Use of distance-ordering generated more consistent accuracy and reliability niches only for the polygon-based methods while the others were affected idiosyncratically. AKDE was disturbed because the ctmm package algorithm did not perform consistently with reordered data generating low sample sizes used to fit GAM (see SI). KDE with distance-ordering did not improve but rather shifted the optimal value of area covered with increasing observations. For MCP distance-ordering only improved the reliability of estimates. The use of distance-ordering allowed better estimates than the raw LoCoh without reaching our criteria. With BicubIt, using distance-ordering did not improve or hampered but changed the shape of the accuracy niche.

### Interacting effects of trap density and area covered

When the number of observations was fixed to 7, depending on the estimator used, only the area covered, or a combination of area covered and trap density values determined the accuracy and reliability. Kernel-based estimators offered reliable estimates the accuracy of which was determined solely by variation in the area covered (Fig. 6, see elongated accuracy niches along the x axis). The patterns for the remaining three estimators were more complex. Only AKDE was able to recover an accurate estimate at intermediate-high values of area covered when estimators were used alone.

**Figure 6:**
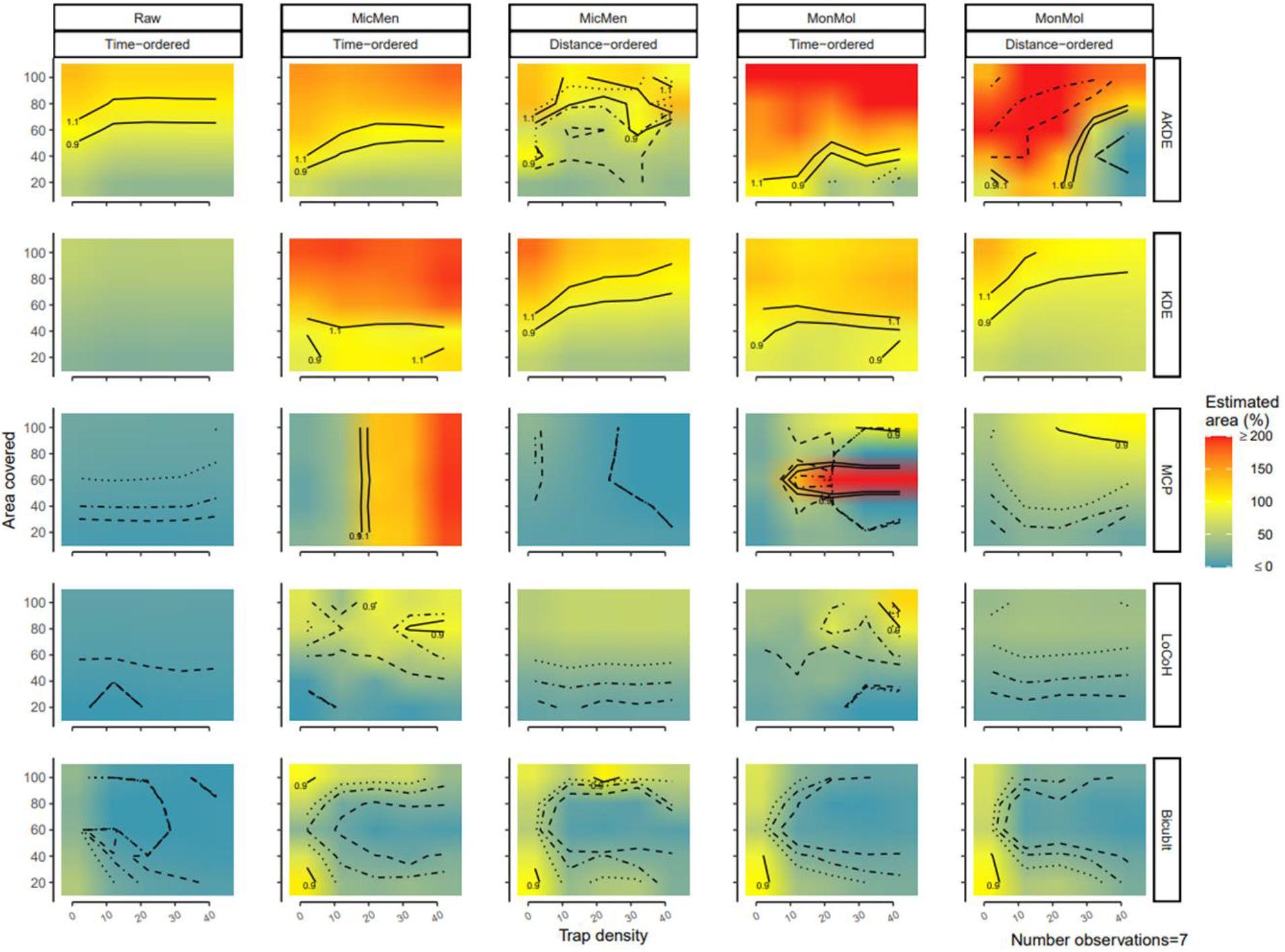
Interacting effects of trap density and area covered by the grid on home range area estimations: Rows gather different estimators, from top: autocorrelated Kernel density estimator (AKDE), the traditional version (KDE), minimum convex polygon (MCP), local convex hull (LoCoH) and bicubic interpolation (Bicublt). Columns gather the combination of two factors, the ordering procedure (time or distance) and the asymptotic model used (“Raw” indicating no model, “MicMen” Michaelis-Menten and “MonMol” Monomolecular). The x axis displays the trap density and the y axis the proportion of the area covered by the trapping grid, the number of observations was fixed at 7. The colors in the plots indicate the estimated area as a percentage of the true area predicted by the GAM model, yellow values approximate the true area, and red indicate over-estimation while blue underestimation. Values above 200% where trimmed and given a 200 while values below 0% where also trimmed as 0 to concentrate color variation on a meaningful range. The solid lines indicate the contour of accurate estimations, defined as ±10% of the true area. Pointed lines indicate the reliable estimations measured as those where the confidence interval was ±20% of the estimated area, point-dashed ±40% and dashed ±60%. Finally, on the lower-right corner the fixed variable value is indicated.

Using asymptotic models yielded improvements in accuracy for all estimators. AKDE improved by reducing the area covered needed for an accurate estimate (MonMol 20% to 40%; MicMen 40% to 60%; for lowest and highest trap densities respectively). KDE greatly benefitted from an increase in accuracy that achieved our criteria when the area covered was between 20 and 60% for all except the lowest and highest trap density values. The use of MonMol allowed accurate estimates at all trap densities but for a narrower range at intermediate area covered values at the lowest trap densities and to 20% at the highest trap densities. Using MicMen allowed MCP to yield reliable and accurate estimates for all area covered values at intermediate trap densities while MonMol only for two values of area covered, 50 and 70%. Asymptotic models helped increasing accuracy of LoCoH but these weren’t reliable according to our criteria. BicubIt was empowered by MicMen yielding accuracy at the smallest trap densities for 20% and 100% area covered while MonMol yielded accurate estimates only at 20% area covered.

Using distance-ordering impacted negatively AKDE impairing its reliability. KDE on the contrary benefited substantially by having more accuracy overall despite needing an increase in area covered (MicMen 40% to 80% and MonMol 60% to 100% for low and medium/large values of trap density respectively). For MCP MonMol with distance-ordering helped retrieving reliable estimates from 40% area covered and accurate at 100% area covered from medium to high values of trap density. LoCoh benefited from distance ordering by increasing reliability, but accuracy was not reached. Distance-ordering did not improve or hamper BicubIt estimates.

## Discussion and conclusion

Our results indicate that estimators used with telemetry and GPS data can yield reliable estimates of home range size when used with trapping data. Depending on the methodology, estimators examined were able to retrieve accurate and reliable estimates even in the challenging scenarios including very low spatial resolution and reduced information per individual. Such performance indicates that, despite having gone underused in favor of more modern methodology, trapping data can continue to prove a valuable source for home range studies.

Under most scenarios investigated, AKDE was able to recover an accurate estimate at lower sample sizes than the other estimators. KDE usually followed behind while MCP, LoCoH and BicubIt needed a much higher number of observations. This behavior is consistent with the accuracy of these estimators on telemetry data (Noonan et al. 2019). The use of asymptotic models helped all estimators except AKDE in reaching an accurate estimation with fewer observations. Nevertheless, counterintuitively, under most scenarios the use of asymptotic models’ predictions reduced the consistency of the reached accuracy. For example, as hypothesized by (Gautestad and Mysterud 1995), MCP (and LoCoH) needed more than 1500 observations to reach the asymptote with the true home range. While using asymptotic models’ predictions helped MCP, LoCoH and BicubIt reduce by one or two orders of magnitude the number observations needed for an accurate estimation, increasing the number of observations further led to changes in the predicted area. The use of distance ordering prior to calculating the home range areas and fitting the asymptotic models allowed both reducing the number of observations albeit more moderately, between 5 to 10 times, while maintaining consistency and attaining reliability at lower sample sizes (see SI). Thus, we suggest the use of asymptotic models in home range analyses cautiously, possibly with distance ordering and additional checks on the reliability of asymptotic estimates as developed in (Haines et al. 2009).

Promisingly, accurate estimates of the home range area were obtained with as few as 3 observations per individual, a finding that could encourage movement ecologists to plan trapping studies and other researchers to exploit their existing trapping datasets for home range estimates. Our findings of accuracy and reliability of estimators for home range analyses with trapping data might offer an additional push to enlarge the range of species and researchers in home range studies. A higher diversity of species and researchers can result in new questions being addressed by a more inclusive research milieu not limited by the large budgets needed for telemetry-based monitoring. A standard metallic foldable trap for small animals might cost on the order of tens of dollars while the average, all-in cost of tracking an animal is on the order of tens of thousands of dollars (Thomas et al. 2011). Since traps can be used for several years if well maintained and are not used for a unique individual, they might turn a smaller budget into a competitive alternative.

Telemetry devices are also best deployed on animals large enough to carry them or else become extremely limited by their small battery life. Although tag sizes have been rapidly decreasing, most species are still too small to be monitored by telemetry (Kays et al. 2015). Our results show how the home range area of small animals unsuited to collaring can be accurately monitored using traps. Nevertheless, trapping in wild animals is not free of costs. Traps might be disease vectors and harmful if not maintained properly. Moreover, the psychophysiological welfare of animals should not be neglected, and the time spent in the trap minimized. Reducing the time in the trap might also improve survival because captured animals are vulnerable to predation. An alternative to live trapping could be the use of camera traps or sound recorders (Chavel et al. 2017). The considerations of trap density, number of observations and area covered all should hold if autocorrelation between detections is taken into account.

Over the range of conditions examined, the proportion of the home range area covered by the trapping grid and the number of observations had a much stronger impact than the trap density. Thus, theoretically, the effort of trapping studies could be redirected to expanding the grid to encompass a greater study area instead of densifying it. However, for animals that have overlapping home ranges, reducing the trap density might lead to reduce the chances of relocating a focal individual several times. Under scarcity of traps other residents may be trapped impeding gathering enough observations per individual. To increase the number of observations per individual and the number of individuals, one could increase the number of traps on each knot of the grid, thereby slightly reducing the working time by clumping traps in space. In addition, in our simulations, traps were highly effective having a range of attraction of 200m for a home range of 1 Ha as we tried to mimic high visibility or bated traps. If traps have a lower range of action, sparsening the grid might significantly reduce the sample size per individual.

Reducing the number of grid knots may also alter other properties in real-world studies. For example, having a sparser grid could lead to more individual home ranges being only partially covered, impacting the information obtained. Depending on the severity of the loss of area covered by the grid, AKDE, KDE or BicubIt should be preferred, the latter two being most performant if used with asymptotic models under the smallest area covered and sparser grids here examined. In combination with monomolecular asymptotic models, BicubIt can retrieve an accurate and reliable estimate at as low as 2-3 observations. Thus, a possibility for reducing the costs of trapping studies would be to drastically sparse the grid and use BicubIt. This could serve for example with trapping studies using sophisticated camera traps. Further investigations on wild populations simultaneously monitored with telemetry or GPS and trapping should be conducted to confirm our findings in such challenging scenarios. We suggest studies reduce the trap density in favor of long term or more intense monitoring but ideally researchers should have a sense of the mean home range radius of the species of interest, the degree of territoriality as well as the range of attraction of the traps. By doing so, they might be able to assess the “sweet spot” between reducing the number of traps and the reliability and accuracy of the information obtained.

In this study we have used the simplest type of home range, one that is circular, and the animal movement independent within it to have a first sense of how common and uncommon methods performed with trapping data. Due to the complexity and computational burden of the range of methods here used we opted for the simplest case of uncorrelated movement detections. Given that the most reliable and accurate estimator was the only one that doesn’t assume uncorrelated observations (AKDE), we suspect that adding autocorrelation to the data shouldn’t change the main conclusions of the study (i.e., that trap data contain sufficient information for home-range estimation). On scenarios with autocorrelated data, the difference between AKDE and the other estimators should broaden as the rest of estimators become impacted by the autocorrelation in the observations. Besides this, animal locations obtained with trapping methods should generally be less autocorrelated, at least for live trapping. The elapsed time between one observation and the next for the same individual should generally exceed the range crossing time. It would be important though in trapping studies to control for the time of the day since animals may repeatedly use the same locations for resting or sleeping. Thus, if traps are always set before the onset of activity, a given individual might be trapped always near the resting place generating autocorrelation. In addition, non-circular shapes of home ranges, which might be common in nature, could trigger biases in estimates (Seaman et al. 1999, Halbrook and Petach 2018). A possibility to improve the presented methodology in this context could be to tune the reference bandwidth allowing non-continuous polygons or use habitat-informed versions of KDE (Halbrook and Petach 2018). Nevertheless, these considerations might be irrelevant for conservation studies were the target might be the area that encompasses all resources used by an animal *sensu* (Burt 1943), even if they use linear corridors to travel between food patches or shelters. Overall, the results of the current study might offer a valuable reference to assess bias in home range analyses with trapping data.

The evidence here presented can act as a useful building block for future research. Although further research might nuance the results here presented, the main finding that trapping data is a useful alternative to more expensive technology for home range size estimates is unlikely to change. The use of trapping data might enlarge the range of ideas that can be explored by broadening the scope of researchers pursuing, and species used in, home range analyses.

## Acknowledgments

We are very grateful to Johannes Signer for his advice on the implementation of functions from the R package “amt”. MJN was supported by an NSERC Discovery Grant RGPIN-2021-02758.

## Authors contributions

L S-M and L P developed the ideas that served as first step of this project. MJ N suggested simulations as a tool for testing, the use of bicubic interpolation and further research goals. L S-M led the coding and wrote the first draft while all authors contributed to coding and writing the final version of the manuscript.

## Data availability

## Figures and tables

Figures were generated using R packages base v 4.0.5 (R Core Team 2019), ggplot2 v 3.3.5 (Wickham 2016), ggpubr 0.4.0 (Kassambara 2020), lemon v 0.4.5 (Edwards 2020) and directlabels v 2021.1.13 (Hocking 2021).

## Notes

### Competing Interest Statement

The authors have declared no competing interest.

## References

Allen, A. M. and Singh, N. J. 2016. Linking Movement Ecology with Wildlife Management and Conservation. - Front. Ecol. Evol. in press.

Andrzejewski, R. 2002. The home-range concept in rodents revised. - Acta Theriol. (Warsz.) 47: 81–101.

Bergstrom, B. J. 1988. Home Ranges of Three Species of Chipmunks (Tamias) as Assessed by Radiotelemetry and Grid Trapping. - J. Mammal. 69: 190–193.

Bolker, B. M. 2008. Ecological models and data in R. - Princeton University Press.

Bondrup-Nielsen, S. 2011. Density estimation as a function of live-trapping grid and home range size. - Can. J. Zool. in press.

Burt, W. H. 1943. Territoriality and home range concepts as applied to mammals. - J. Mammal. 24: 346–352.

Chavel, E. E. et al. 2017. Comparative evaluation of three sampling methods to estimate detection probability of American red squirrels (Tamiasciurus hudsonicus). - Mamm. Biol. 83: 1–9.

Dammhahn, M. and Kappeler, P. M. 2005. Social system of Microcebus berthae, the world’s smallest primate. - Int. J. Primatol. 26: 407–435.

Dammhahn, M. and Kappeler, P. M. 2008. Small-scale coexistence of two mouse lemur species (Microcebus berthae and M. murinus) within a homogeneous competitive environment. - Oecologia 157: 473–483.

Ebersole, J. P. 1980. Food density and territory size: an alternative model and a test on the reef fish Eupomacentrus leucostictus. - Am. Nat. 115: 492–509.

Edwards, S. M. 2020. lemon: Freshing Up your “ggplot2” Plots.

Fleming, C. H. and Calabrese, J. M. 2017. A new kernel density estimator for accurate home-range and species-range area estimation. - Methods Ecol. Evol. 8: 571–579.

Fleming, C. H. et al. 2014. From Fine-Scale Foraging to Home Ranges: A Semivariance Approach to Identifying Movement Modes across Spatiotemporal Scales. - Am. Nat. 183: E154–E167.

Fleming, C. H. et al. 2018. Correcting for missing and irregular data in home-range estimation. - Ecol. Appl. 28: 1003–1010.

Fleming, C. H. et al. 2019. Overcoming the challenge of small effective sample sizes in home-range estimation. - Methods Ecol. Evol. in press.

Gautestad, A. O. and Mysterud, I. 1995. The Home Range Ghost. - Oikos 74: 195–204.

Gil-Sánchez, J. M. et al. 2011. The use of camera trapping for estimating Iberian lynx (Lynx pardinus) home ranges. - Eur. J. Wildl. Res. 57: 1203–1211.

Haines, A. et al. 2009. A Method for Determining Asymptotes of Home-Range Area Curves. - Natl. Quail Symp. Proc. in press.

Halbrook, R. S. and Petach, M. 2018. Estimated mink home ranges using various home-range estimators. - Wildl. Soc. Bull. 42: 656–666.

Harris, S. et al. 1990. Home-range analysis using radio-tracking data–a review of problems and techniques particularly as applied to the study of mammals. - Mammal Rev. 20: 97–123.

Hayne, D. W. 1949. Calculation of Size of Home Range. - J. Mammal. 30: 1–18.

Hayne, D. W. 1950. Apparent Home Range of Microtus in Relation to Distance between Traps. - J. Mammal. 31: 26–39.

Heupel, M. R. et al. 2004. Estimation of Shark Home Ranges using Passive Monitoring Techniques. - Environ. Biol. Fishes 71: 135–142.

Hocking, T. D. 2021. directlabels: Direct Labels for Multicolor Plots.

Innes, J. G. and Skipworth, J. P. 1983. Home ranges of ship rats in a small New Zealand forest as revealed by trapping and tracking. - N. Z. J. Zool. 10: 99–110.

Kane, M. D. et al. 2015. Potential for camera-traps and spatial mark-resight models to improve monitoring of the critically endangered West African lion (Panthera leo). - Biodivers. Conserv. 24: 3527–3541.

Kassambara, A. 2020. ggpubr: “ggplot2” Based Publication Ready Plots.

Kays, R. et al. 2015. Terrestrial animal tracking as an eye on life and planet. - Science 348: aaa2478.

Kie, J. G. 2013. A rule-based ad hoc method for selecting a bandwidth in kernel home-range analyses. - Anim. Biotelemetry 1: 13.

Kie, J. G. et al. 2010. The home-range concept: are traditional estimators still relevant with modern telemetry technology? - Philos. Trans. R. Soc. Lond. B. Biol. Sci. 365: 2221–2231.

Kumbhojkar, S. et al. 2020. A Camera-Trap Home-Range Analysis of the Indian Leopard (Panthera pardus fusca) in Jaipur, India. - Animals 10: 1600.

Laver, P. N. and Kelly, M. J. 2008. A critical review of home range studies. - J. Wildl. Manag. 72: 290–298.

Law, B. S. and Dickman, C. R. 1998. The use of habitat mosaics by terrestrial vertebrate fauna: implications for conservation and management. - Biodivers. Conserv. 7: 323–333.

Leo, B. T. et al. 2016. Home Range Estimates of Feral Cats (Felis catus) on Rota Island and Determining Asymptotic Convergence1. - Pac. Sci. 70: 323–331.

Lira, P. K. and Fernandez, F.A. dos S. 2009. A comparison of trapping- and radiotelemetry-based estimates of home range of the neotropical opossum Philander frenatus. - Mamm. Biol. 74: 1–8.

Lukacs, P. M. et al. 2005. Evaluation of trapping-web designs. - Wildl. Res. 32: 103–110.

Mohr, C. O. 1947. Table of Equivalent Populations of North American Small Mammals. - Am. Midl. Nat. 37: 223–249.

Morato, R. G. et al. 2016. Space Use and Movement of a Neotropical Top Predator: The Endangered Jaguar (M Stöck, Ed.). - PLOS ONE 11: e0168176.

Mueller, T. and Fagan, W. F. 2008. Search and navigation in dynamic environments–from individual behaviors to population distributions. - Oikos 117: 654–664.

Nathan, R. et al. 2008. A movement ecology paradigm for unifying organismal movement research. - Proc. Natl. Acad. Sci. 105: 19052–19059.

Noonan, M. J. et al. 2019. A comprehensive analysis of autocorrelation and bias in home range estimation. - Ecol. Monogr. 89: e01344.

Oppel, S. et al. 2018. Spatial scales of marine conservation management for breeding seabirds. - Mar. Policy 98: 37–46.

Ouellette, M. and Cardille, J. A. 2011. The Complex Linear Home Range Estimator: Representing the Home Range of River Turtles Moving in Multiple Channels. - Chelonian Conserv. Biol. 10: 259–265.

Plotz, R. D. et al. 2016. Standardising Home Range Studies for Improved Management of the Critically Endangered Black Rhinoceros. - PloS One 11: e0150571.

Powell, R. A. and Mitchell, M. S. 2012. What is a home range? - J. Mammal. 93: 948–958.

Powell, R. A. et al. 1997. Ecology and behaviour of North American black bears: home ranges, habitat, and social organization. - Chapman & Hall.

R Core Team 2019. R: A Language and Environment for Statistical Computing. - R Foundation for Statistical Computing.

Rajska-Jurgiel, E. 2001. Movement behaviour of woodland rodents: looking from beyond small trapping grids. - Acta Theriol. (Warsz.) 46: 145–159.

Reid, M. L. and Weatherhead, P. J. 1988. Topographical constraints on competition for territories. - Oikos: 115–117.

Ribble, D. O. et al. 2002. A Comparison of Home Ranges of two Species of Peromyscus Using Trapping and Radiotelemetry Data. - J. Mammal. 83: 260–266.

Rose, B. 1982. Lizard Home Ranges: Methodology and Functions. - J. Herpetol. 16: 253–269.

Schick, R. S. et al. 2008. Understanding movement data and movement processes: current and emerging directions. - Ecol. Lett. 11: 1338–1350.

Schoener, T. W. 1981. An empirically based estimate of home range. - Theor. Popul. Biol. 20: 281–325.

Seaman, D. E. et al. 1999. Effects of Sample Size on Kernel Home Range Estimates. - J. Wildl. Manag. 63: 739–747.

Silva, I. et al. 2022. Autocorrelation-informed home range estimation: A review and practical guide. - Methods Ecol. Evol. 13: 534–544.

Soanes, L. M. et al. 2013. How many seabirds do we need to track to define home-range area? - J. Appl. Ecol. 50: 671–679.

Spencer, S. R. et al. 1990. Operationally Defining Home Range: Temporal Dependence Exhibited by Hispid Cotton Rats. - Ecology 71: 1817–1822.

Sun, C. C. et al. 2014. Trap Configuration and Spacing Influences Parameter Estimates in Spatial Capture- Recapture Models. - PLOS ONE 9: e88025.

Thomas, B. et al. 2011. Wildlife tracking technology options and cost considerations. - Wildl. Res. 38: 653–663.

Van Winkle, W. 1975. Comparison of Several Probabilistic Home-Range Models. - J. Wildl. Manag. 39: 118–123.

Vieira, W. F. et al. 2019. A comparison of methods to determine chimpanzee home-range size in a forest-farm mosaic at Madina in Cantanhez National Park, Guinea-Bissau. - Primates J. Primatol. 60: 355–365.

Ward, G. D. 1984. Comparison of trap- and radio-revealed home ranges of the brush-tailed possum (Trichosurus vulpecula Kerr) in New Zealand lowland forest. - N. Z. J. Zool. 11: 85–92.

Wickham, H. 2016. ggplot2: Elegant Graphics for Data Analysis. - Springer-Verlag New York.

Wood, S. N. 2017. Generalized Additive Models: An Introduction with R, Second Edition. - Taylor & Francis Inc.

Worton, B. J. 1987. A review of models of home range for animal movement. - Ecol. Model. 38: 277–298.

Worton, B. J. 1989. Kernel methods for estimating the utilization distribution in home-range studies. - Ecology 70: 164–168.

Wszola, L. S. et al. 2019. Simulating detection-censored movement records for home range analysis planning. - Ecol. Model. 392: 268–278.

