## Supplementary information for "Are trapping data still suited for home range estimation? An analysis with various estimators, asymptotic models and data ordering procedures"

### Generalized additive model

#### Formula

Cubic regression spline was chosen for the number of observations variable in order to be able to place the knots manually (see main text).

```
#type of spline

spl<-c("cr","tp","tp")

knbobs<-15 #number of knots

form.ti<-as.formula(area~

        intsem

       +s(log10(nbobs),k=knbobs,bs=spl[1],by=intsem)

       +s(trap.d,k=5,bs=spl[2],by=intsem)

       +s(area.c,k=5,bs=spl[3],by=intsem)

       +ti(log10(nbobs),trap.d,k=c(knbobs,5),bs=spl[c(1,2)],by=intsem)

       +ti(log10(nbobs),area.c,k=c(knbobs,5),bs=spl[c(1,3)],by=intsem)

       +ti(trap.d,area.c,k=c(5,5),bs=spl[c(2,3)],by=intsem)

       +ti(log10(nbobs),area.c,trap.d,k=c(knbobs,5,5),bs=spl[c(1,2,3)],by=intsem)

)
```

#### Diagnostics

Residuals have a mean close to 0 (see SI figures 1, 2 and 4) and most are close to the mean (SI Fig. 4), indicating that the predictions from the GAM model are in accordance with the data from the simulations. Nevertheless, there is unequal variance, and residuals are not normally distributed. These violations of assumptions might be present for two reasons: 1. The response variable “area” is a proportion bounded at 0 but potentially unbounded towards positive infinity. At low sample sizes, the home range size estimates might change quickly because a new observation thus exerting a disproportionate impact on the estimated asymptotes (see SI Fig. 2). Such asymptotes sometimes overestimate the true area by several orders of magnitude, thus generating outliers towards the positive range of the residuals. 2. The presence of right skew on the positive side of the residuals might also be a consequence that we sampled more especially the “problematic” regions, the low end of the number of observations. Interestingly, although this problem is especially relevant for asymptotic model predictions before ~60 observations are gathered (SI Fig. 2), the use of distance-ordering restrains the high variance zone to a much lower range (~10 KDE, ~20 MCP and LoCoH). The other estimators either do not show unequal variance (BicubIt) or have a better equal low variance used with raw data (AKDE after ~7 observations). Overall, these violations might not impact the conclusions of the study, because i) the vast majority of residuals are close to 0 (SI Fig. 4), and the results of the analyses conducted already indicate that ii) AKDE is the most performant estimator and that iii) distance-ordering is allows better consistency and accuracy for the other estimators.

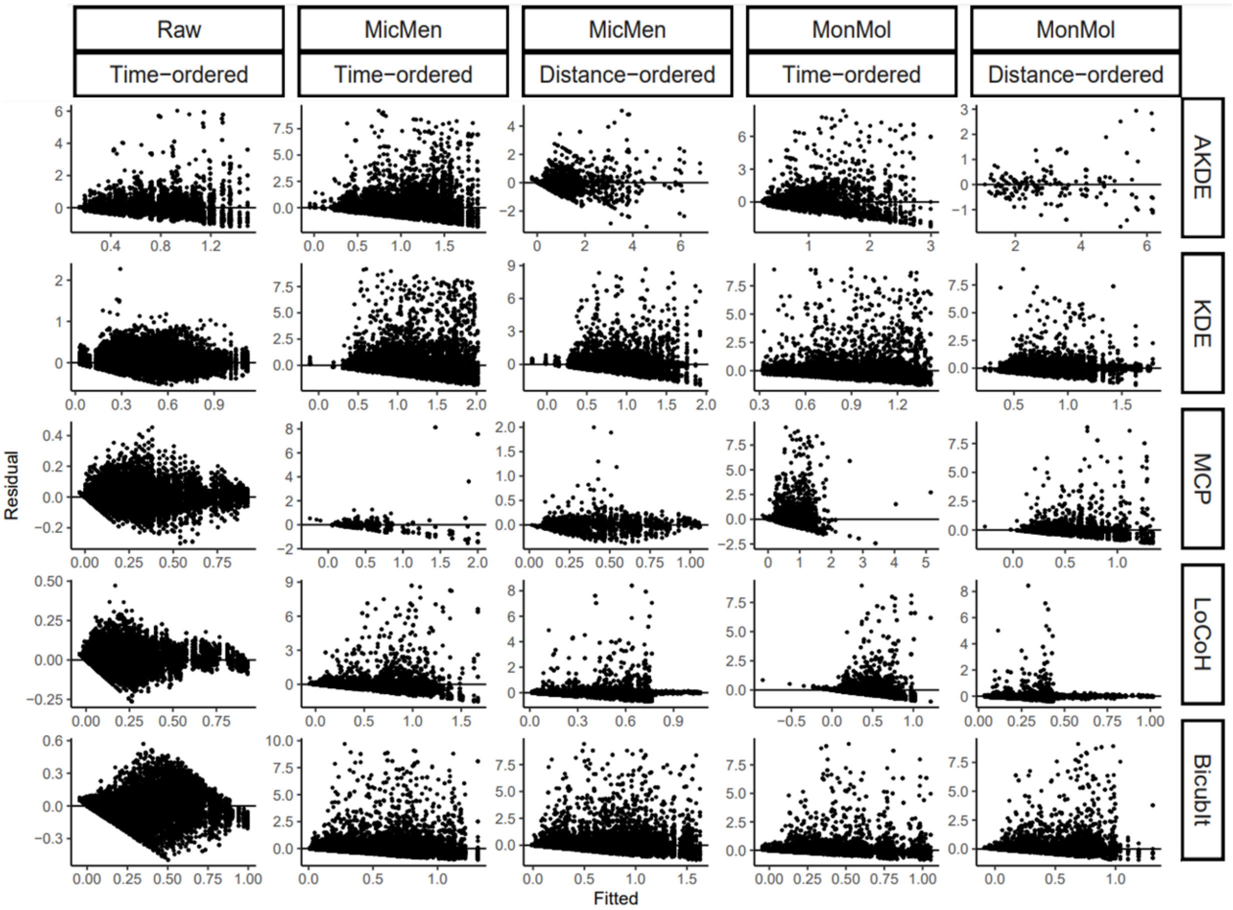


**SI Figure 1: GAM model residuals and fitted values:** Rows gather different estimators, from top: autocorrelated Kernel density estimator (AKDE), the traditional version (KDE), minimum convex polygon (MCP), local convex hull (LoCoH) and bicubic interpolation (BicubIt). Columns gather the combination of two factors, the ordering procedure (time or distance) and the asymptotic model used ("Raw" indicating no model, "MicMen" Michaelis-Menten and "MonMol" Monomolecular). The x axis displays the fitted model values and the y axis the corresponding residuals.

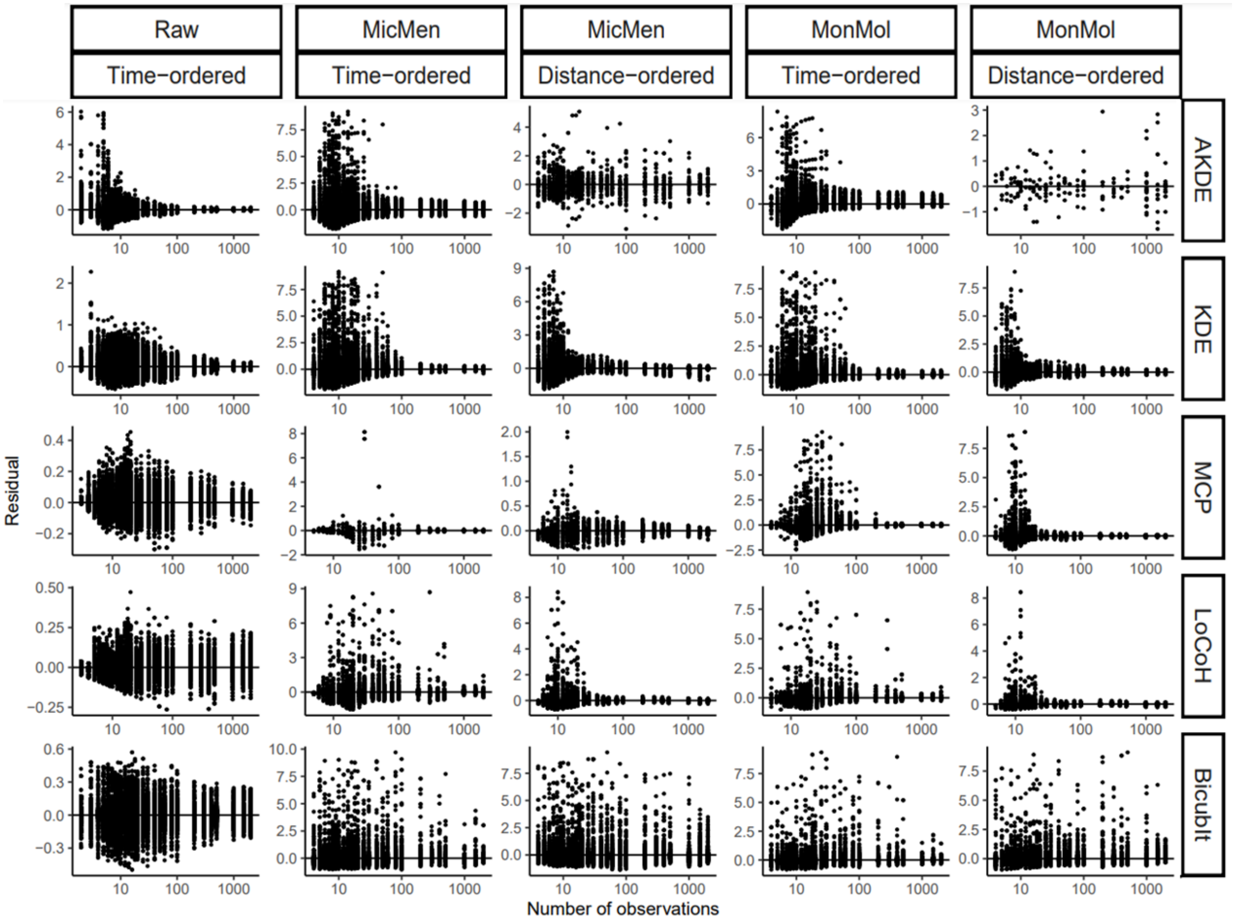


**SI Figure 2: GAM model residuals conditional on the number of observations:** Rows gather different estimators, from top: autocorrelated Kernel density estimator (AKDE), the traditional version (KDE), minimum convex polygon (MCP), local convex hull (LoCoH) and bicubic interpolation (BicubIt). Columns gather the combination of two factors, the ordering procedure (time or distance) and the asymptotic model used ("Raw" indicating no model, "MicMen" Michaelis-Menten and "MonMol" Monomolecular). The x axis displays the number of observations values and the y axis the corresponding residuals.

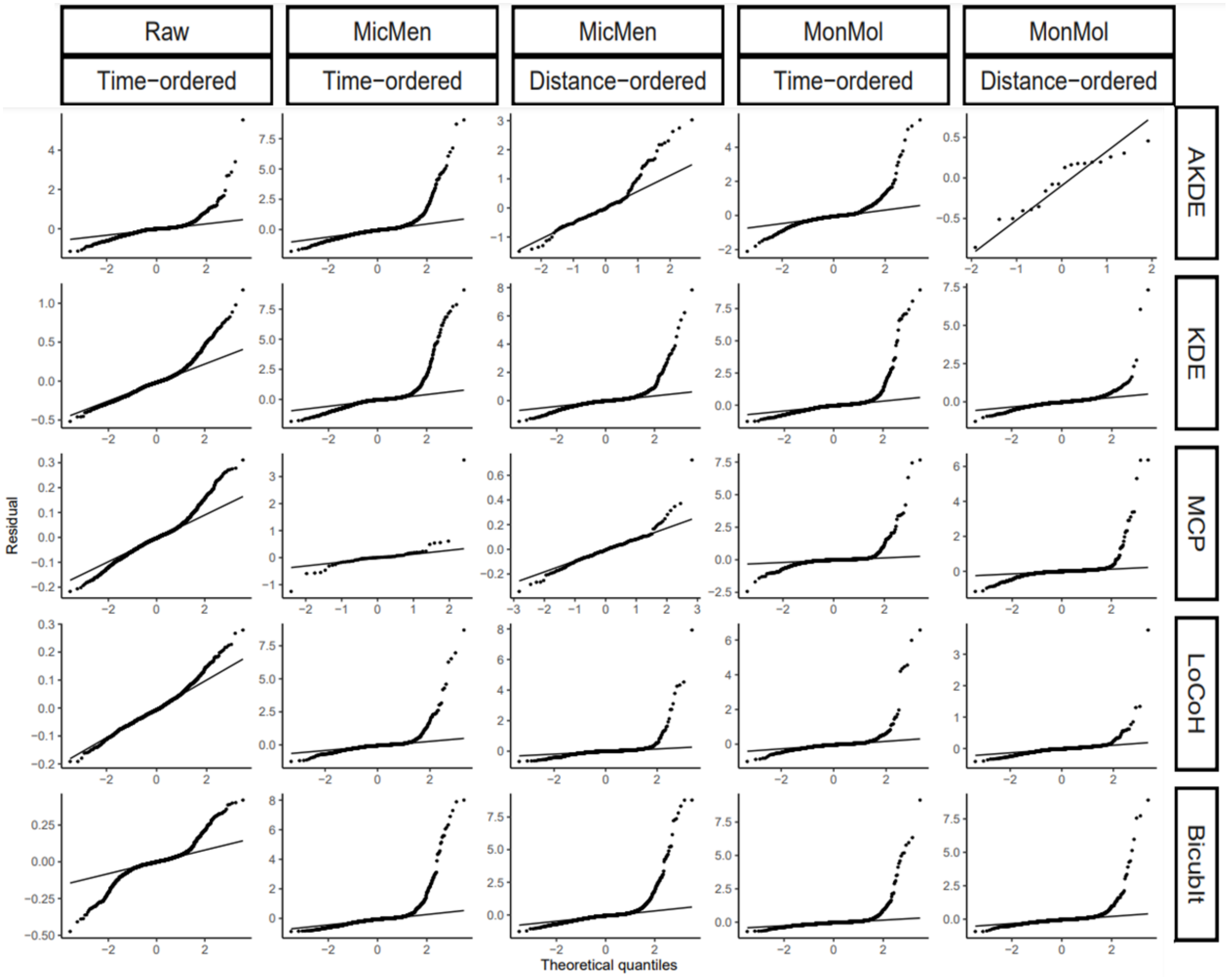


**SI Figure 3: Quantile-quantile plot of GAM model residuals:** Rows gather different estimators, from top: autocorrelated Kernel density estimator (AKDE), the traditional version (KDE), minimum convex polygon (MCP), local convex hull (LoCoH) and bicubic interpolation (BicubIt). Columns gather the combination of two factors, the ordering procedure (time or distance) and the asymptotic model used ("Raw" indicating no model, "MicMen" Michaelis-Menten and "MonMol" Monomolecular). The x axis displays the theoretical quantiles and the y axis the corresponding residuals.

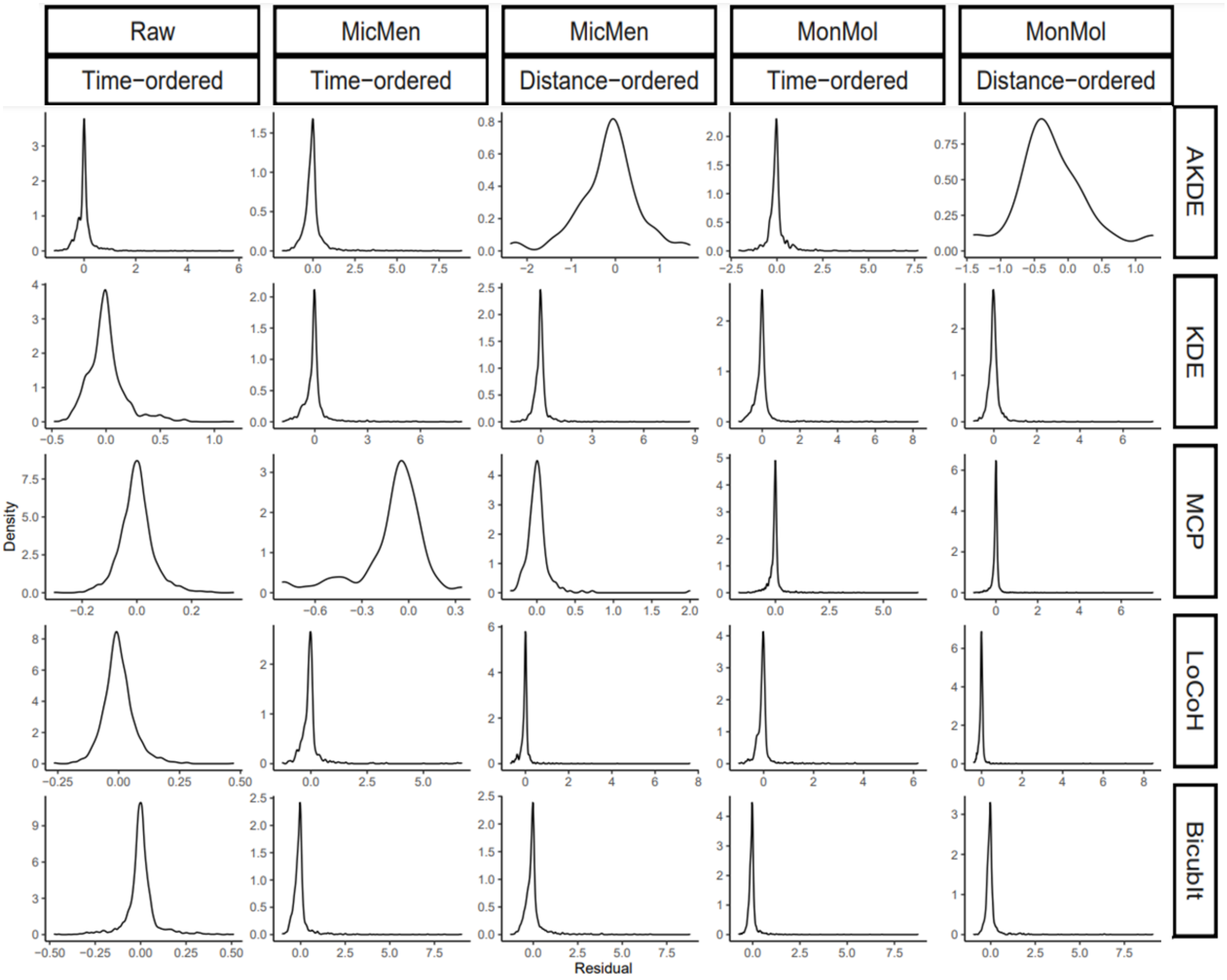


**SI Figure 4: Density plot of GAM model residuals:** Rows gather different estimators, from top: autocorrelated Kernel density estimator (AKDE), the traditional version (KDE), minimum convex polygon (MCP), local convex hull (LoCoH) and bicubic interpolation (BicubIt). Columns gather the combination of two factors, the ordering procedure (time or distance) and the asymptotic model used ("Raw" indicating no model, "MicMen" Michaelis-Menten and "MonMol" Monomolecular). The x axis displays the residuals and the y axis the corresponding density.

### Additional plots

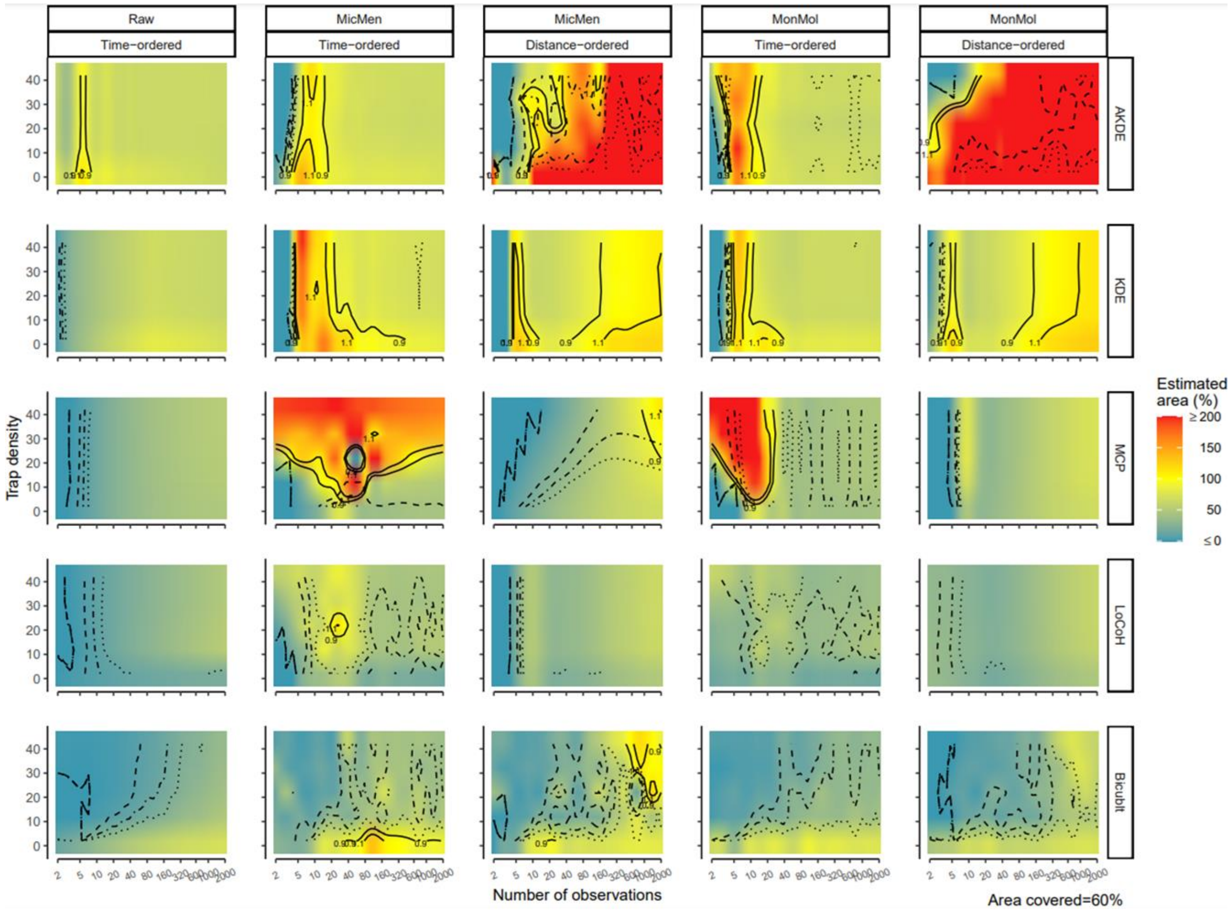


**SI Figure 5: Interacting effects of number of observations and trap density on home range area estimations:** Rows gather different estimators, from top: autocorrelated Kernel density estimator (AKDE), the traditional version (KDE), minimum convex polygon (MCP), local convex hull (LoCoH) and bicubic interpolation (BicubIt). Columns gather the combination of two factors, the ordering procedure (time or distance) and the asymptotic model used ("Raw" indicating no model, "MicMen" Michaelis-Menten and "MonMol" Monomolecular). The x axis displays the number of observations and the y axis the trap density. The colors in the plots indicate the estimated area as a percentage of the true area predicted by the GAM model, yellow values approximate the true area, and red indicate over-estimation while blue underestimation. Values above 200% where trimmed and given a 200 while values below 0% where also trimmed as 0 to concentrate color variation on a meaningful range. The solid lines indicate the contour of accurate estimations, defined as ±10% of the true area. Pointed lines indicate the reliable estimations measured as those where the confidence interval was ±20% of the estimated area, point-dashed ±40% and dashed ±60%. Finally, on the lower-right corner the fixed variable value is indicated.

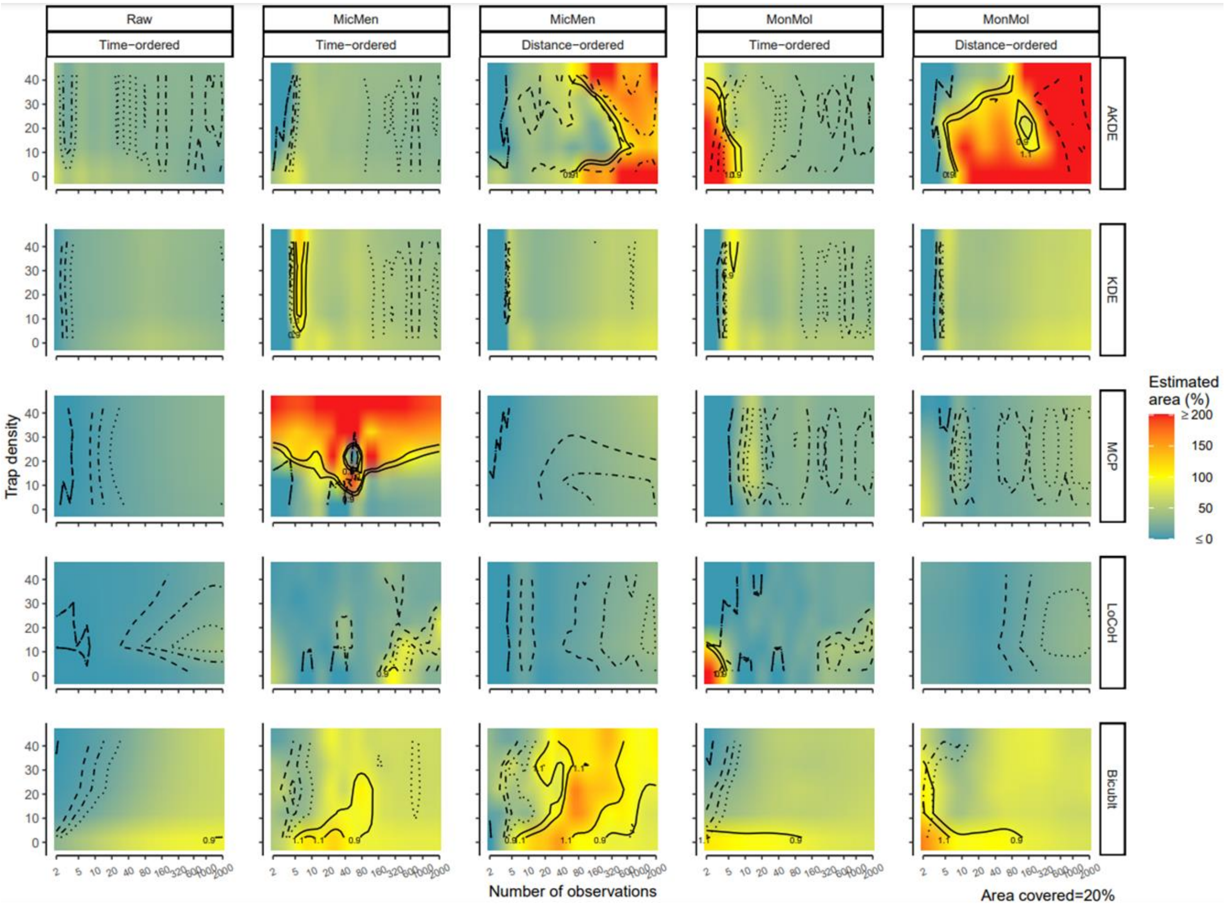


**SI Figure 6: Interacting effects of number of observations and trap density on home range area estimations:** Rows gather different estimators, from top: autocorrelated Kernel density estimator (AKDE), the traditional version (KDE), minimum convex polygon (MCP), local convex hull (LoCoH) and bicubic interpolation (BicubIt). Columns gather the combination of two factors, the ordering procedure (time or distance) and the asymptotic model used ("Raw" indicating no model, "MicMen" Michaelis-Menten and "MonMol" Monomolecular). The x axis displays the number of observations and the y axis the trap density. The colors in the plots indicate the estimated area as a percentage of the true area predicted by the GAM model, yellow values approximate the true area, and red indicate over-estimation while blue underestimation. Values above 200% where trimmed and given a 200 while values below 0% where also trimmed as 0 to concentrate color variation on a meaningful range. The solid lines indicate the contour of accurate estimations, defined as ±10% of the true area. Pointed lines indicate the reliable estimations measured as those where the confidence interval was ±20% of the estimated area, point-dashed ±40% and dashed ±60%. Finally, on the lower-right corner the fixed variable value is indicated.

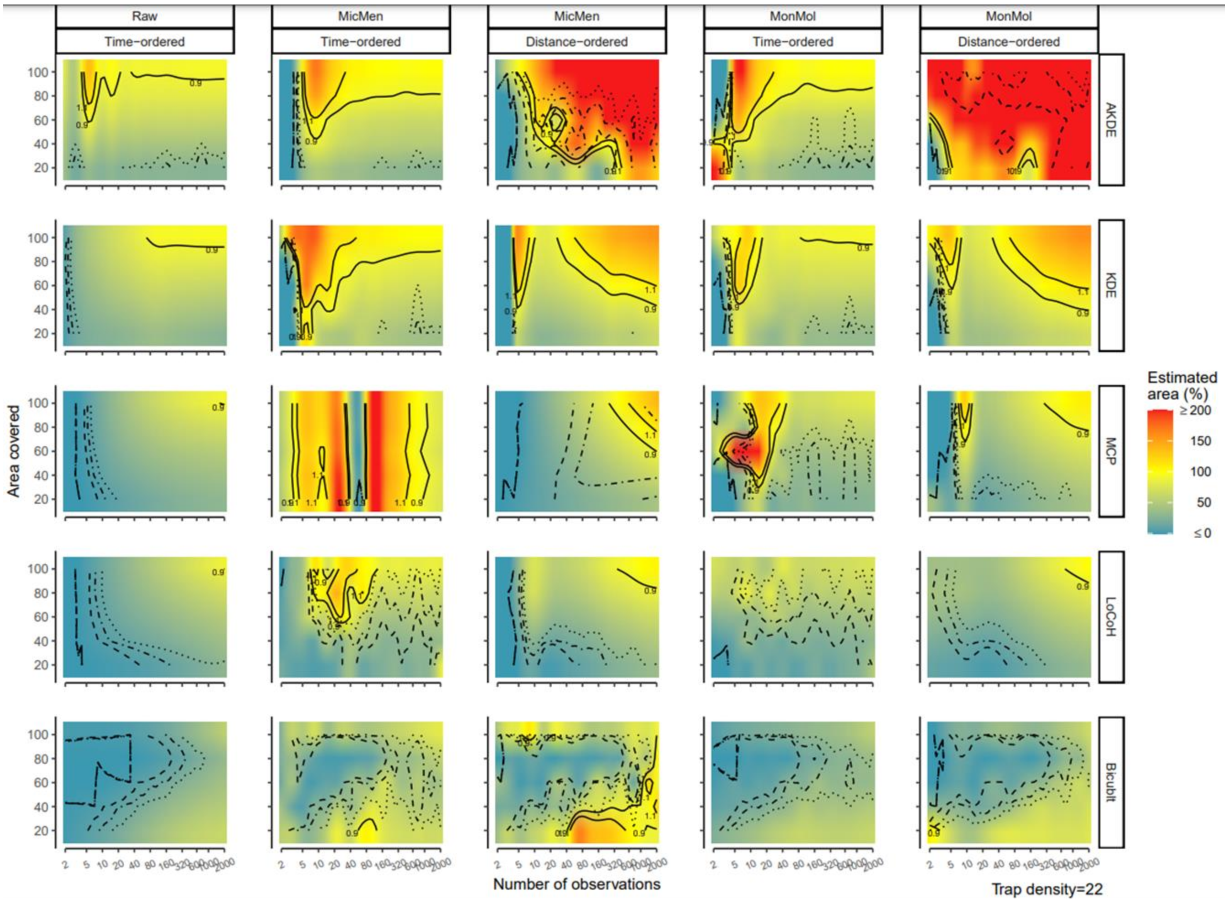


**SI Figure 7: Interacting effects of number of observations and area covered by the grid on home range area estimations:** Rows gather different estimators, from top: autocorrelated Kernel density estimator (AKDE), the traditional version (KDE), minimum convex polygon (MCP), local convex hull (LoCoH) and bicubic interpolation (BicubIt). Columns gather the combination of two factors, the ordering procedure (time or distance) and the asymptotic model used ("Raw" indicating no model, "MicMen" Michaelis-Menten and "MonMol" Monomolecular). The x axis displays the number of observations and the y axis the proportion of the area covered by the trapping grid. The colors in the plots indicate the estimated area as a percentage of the true area predicted by the GAM model, yellow values approximate the true area, and red indicate over-estimation while blue underestimation. Values above 200% where trimmed and given a 200 while values below 0% where also trimmed as 0 to concentrate color variation on a meaningful range. The solid lines indicate the contour of accurate estimations, defined as ±10% of the true area. Pointed lines indicate the reliable estimations measured as those where the confidence interval was ±20% of the estimated area, point-dashed ±40% and dashed ±60%. Finally, on the lower-right corner the fixed variable value is indicated.

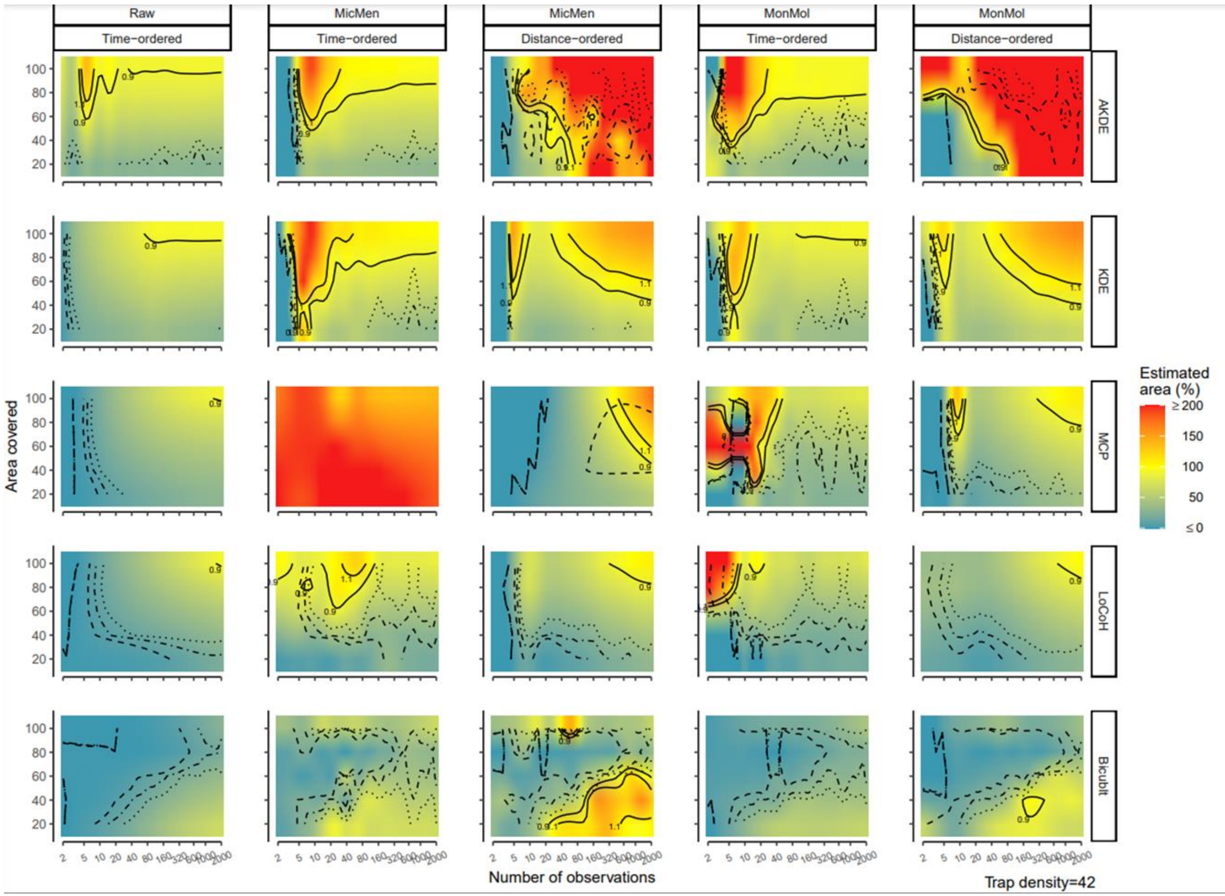


**SI Figure 8: Interacting effects of number of observations and area covered by the grid on home range area estimations:** Rows gather different estimators, from top: autocorrelated Kernel density estimator (AKDE), the traditional version (KDE), minimum convex polygon (MCP), local convex hull (LoCoH) and bicubic interpolation (BicubIt). Columns gather the combination of two factors, the ordering procedure (time or distance) and the asymptotic model used ("Raw" indicating no model, "MicMen" Michaelis-Menten and "MonMol" Monomolecular). The x axis displays the number of observations and the y axis the proportion of the area covered by the trapping grid. The colors in the plots indicate the estimated area as a percentage of the true area predicted by the GAM model, yellow values approximate the true area, and red indicate over-estimation while blue underestimation. Values above 200% where trimmed and given a 200 while values below 0% where also trimmed as 0 to concentrate color variation on a meaningful range. The solid lines indicate the contour of accurate estimations, defined as ±10% of the true area. Pointed lines indicate the reliable estimations measured as those where the confidence interval was ±20% of the estimated area, point-dashed ±40% and dashed ±60%. Finally, on the lower-right corner the fixed variable value is indicated.

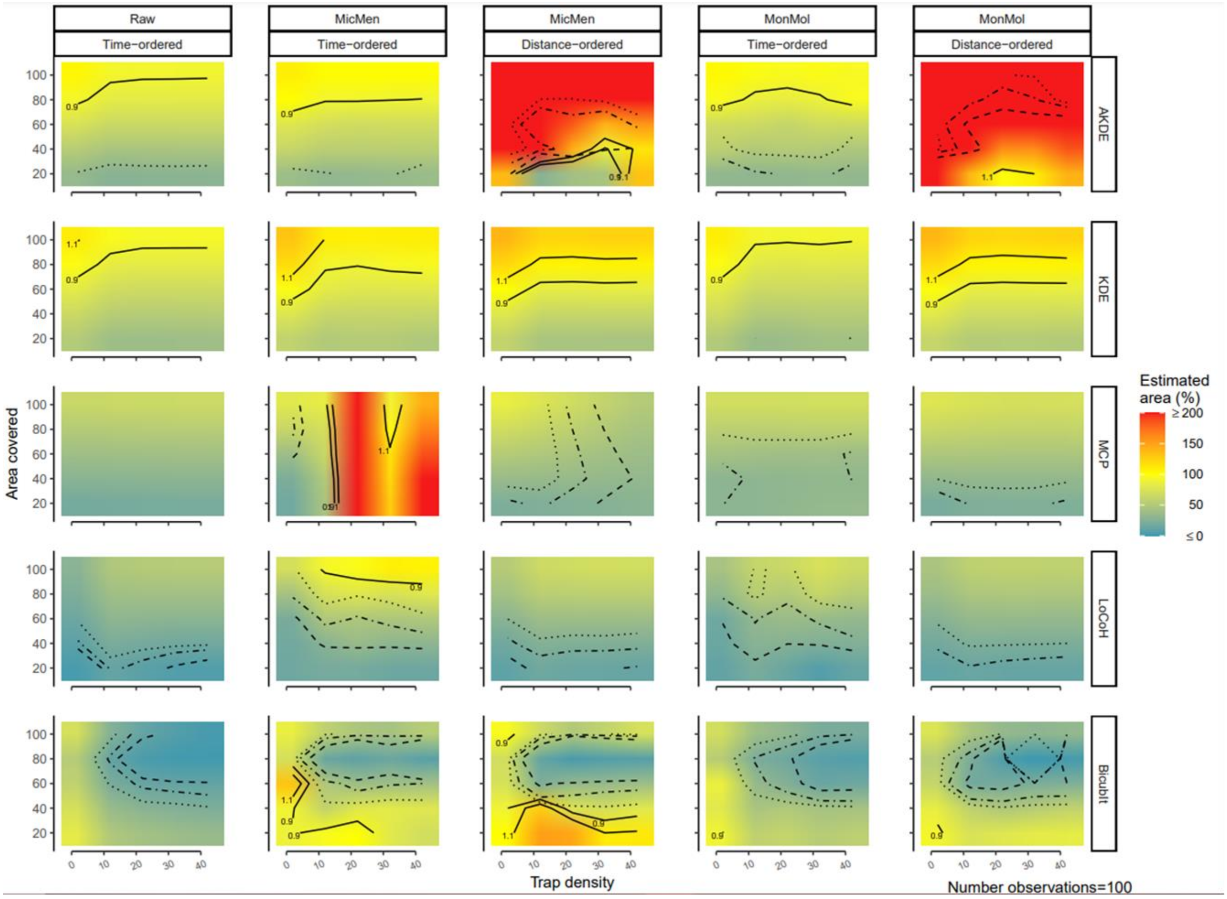


**SI Figure 9: Interacting effects of trap density and area covered by the grid on home range area estimations:** Rows gather different estimators, from top: autocorrelated Kernel density estimator (AKDE), the traditional version (KDE), minimum convex polygon (MCP), local convex hull (LoCoH) and bicubic interpolation (BicubIt). Columns gather the combination of two factors, the ordering procedure (time or distance) and the asymptotic model used ("Raw" indicating no model, "MicMen" Michaelis-Menten and "MonMol" Monomolecular). The x axis displays the trap density and the y axis the proportion of the area covered by the trapping grid. The colors in the plots indicate the estimated area as a percentage of the true area predicted by the GAM model, yellow values approximate the true area, and red indicate over-estimation while blue underestimation. Values above 200% where trimmed and given a 200 while values below 0% where also trimmed as 0 to concentrate color variation on a meaningful range. The solid lines indicate the contour of accurate estimations, defined as ±10% of the true area. Pointed lines indicate the reliable estimations measured as those where the confidence interval was ±20% of the estimated area, point-dashed ±40% and dashed ±60%. Finally, on the lower-right corner the fixed variable value is indicated.

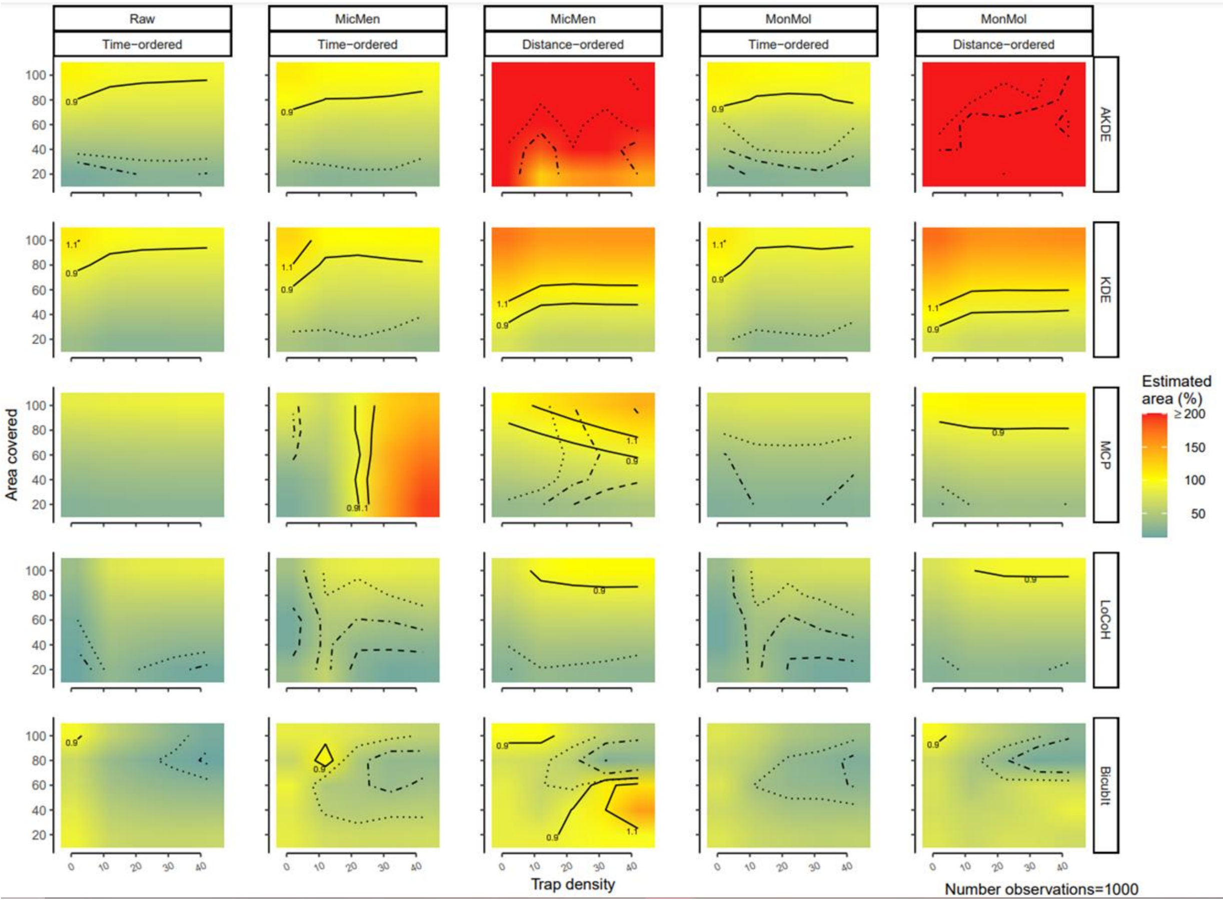


**SI Figure 10: Interacting effects of trap density and area covered by the grid on home range area estimations:** Rows gather different estimators, from top: autocorrelated Kernel density estimator (AKDE), the traditional version (KDE), minimum convex polygon (MCP), local convex hull (LoCoH) and bicubic interpolation (BicubIt). Columns gather the combination of two factors, the ordering procedure (time or distance) and the asymptotic model used ("Raw" indicating no model, "MicMen" Michaelis-Menten and "MonMol" Monomolecular). The x axis displays the trap density and the y axis the proportion of the area covered by the trapping grid. The colors in the plots indicate the estimated area as a percentage of the true area predicted by the GAM model, yellow values approximate the true area, and red indicate over-estimation while blue underestimation. Values above 200% where trimmed and given a 200 while values below 0% where also trimmed as 0 to concentrate color variation on a meaningful range. The solid lines indicate the contour of accurate estimations, defined as ±10% of the true area. Pointed lines indicate the reliable estimations measured as those where the confidence interval was ±20% of the estimated area, point-dashed ±40% and dashed ±60%. Finally, on the lower-right corner the fixed variable value is indicated.

### Limitation of Kernel based methods with increasing number of observations

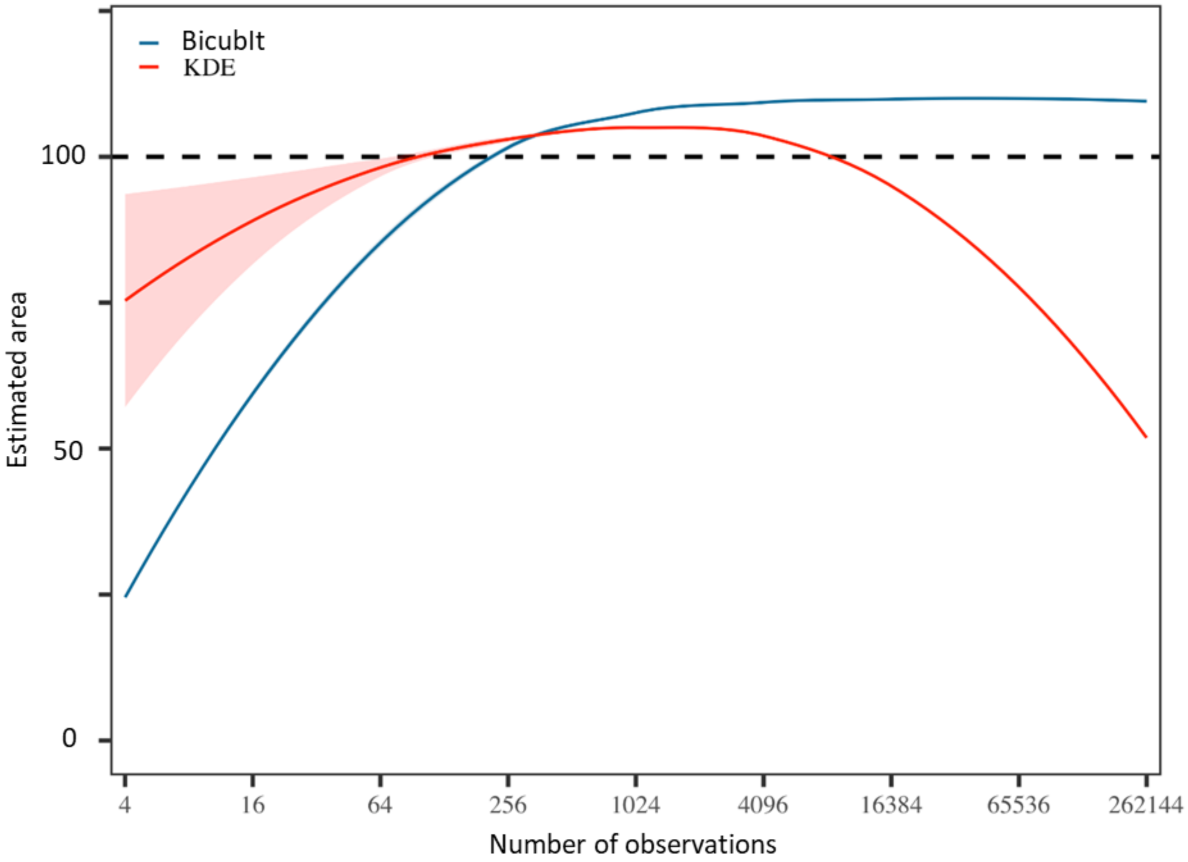


**SI Figure 11: Asymptotic consistency under very high sample sizes:** The traditional Kernel density estimation (KDE, shown in red) estimated areas are compared to those obtained with Bicubic interpolation (BicubIt, blue). Shaded areas correspond to 95% confidence interval. The x axis shows the number of observations, while the y axis shows the percentage of the true area in size recovered by the given estimate.
